# Differential relationships between brain structure and dual task walking in young and older adults

**DOI:** 10.1101/2021.11.04.467303

**Authors:** KE Hupfeld, JM Geraghty, HR McGregor, CJ Hass, O Pasternak, RD Seidler

## Abstract

Almost 25% of all older adults experience difficulty walking. Mobility difficulties for older adults are more pronounced when performing a simultaneous cognitive task while walking (i.e., dual task walking). Although it is known that aging results in widespread brain atrophy, few studies have integrated across more than one neuroimaging modality to comprehensively examine the structural neural correlates that may underly dual task walking in older age. We collected spatiotemporal gait data during single and dual task walking for 37 young (18-34 years) and 23 older adults (66-86 years). We also collected *T*_1_-weighted and diffusion-weighted MRI scans to determine how brain structure differs in older age and relates to dual task walking. We addressed two aims: 1) to characterize age differences in brain structure across a range of metrics including volumetric, surface, and white matter microstructure; and 2) to test for age group differences in the relationship between brain structure and the dual task cost (DTcost) of gait speed and variability. Key findings included widespread brain atrophy for the older adults, with the most pronounced age differences in brain regions related to sensorimotor processing. We also found multiple associations between regional brain atrophy and greater DTcost of gait speed and variability for the older adults. The older adults showed a relationship of both thinner temporal cortex and shallower sulcal depth in the frontal, sensorimotor, and parietal cortices with greater DTcost of gait. Additionally, the older adults showed a relationship of ventricular volume and superior longitudinal fasciculus free-water corrected axial and radial diffusivity with greater DTcost of gait. These relationships were not present for the young adults. Stepwise multiple regression found sulcal depth in the left precentral gyrus, axial diffusivity in the superior longitudinal fasciculus, and sex to best predict DTcost of gait speed, and cortical thickness in the superior temporal gyrus to best predict DTcost of gait variability for older adults. These results contribute to scientific understanding of how individual variations in brain structure are associated with mobility function in aging. This has implications for uncovering mechanisms of brain aging and for identifying target regions for mobility interventions for aging populations.

**CONTRIBUTION TO THE FIELD:** Older age is associated with poorer mobility, including difficulties in performing a cognitive task while walking (i.e., dual task walking). Our work contributes to the field by examining multimodal structural neuroimaging data to characterize how brain structure relates to dual task walking in young versus older adults. We extracted multiple indices from *T*_1_-weighted and diffusion-weighted magnetic resonance imaging (MRI) scans that describe morphological characteristics of brain gray matter, white matter, and cerebrospinal fluid. We analyzed MRI and gait data from 37 young (18-34 years) and 23 older (66-86 years) adults. We identified multiple relationships between regional brain atrophy and greater dual task costs (DTcosts) to gait, i.e., greater slowing of gait speed and greater increases in gait variability from single to dual task walking. Specifically, for the older adults only, thinner temporal cortex and shallower sulcal depth in the frontal, sensorimotor, and parietal cortices were associated with larger DTcosts to walking. Additionally, for the older adults only, ventricular volume and superior longitudinal fasciculus free-water corrected axial and radial diffusivity were associated with larger DTcosts. These findings illustrate that temporal, frontoparietal and sensorimotor brain structures are associated with walking DTcosts in older adults, highlighting potential targets for interventions.

## 1 INTRODUCTION

Nearly 25 percent of older adults report serious mobility problems such as difficulty walking or climbing stairs (Kraus, 2016). Older adults tend to encounter even greater difficulty with performing a secondary cognitive task while walking, i.e., dual task walking (e.g., Hollman et al. 2007; Malcolm et al. 2015; Smith et al. 2016; Springer et al. 2006). A common measure of dual task walking performance is dual task cost (DTcost), or the magnitude of performance decline when conducting two tasks at once as opposed to individually (Bayot et al.,2020; Yogev-Seligmann et al., 2008). Older adults typically exhibit greater DTcosts compared with young adults, such as greater slowing of gait speed during dual conditions (for review, see Al-Yahya et al., 2011; Beurskens and Bock, 2012). Examining DTcost is considered more useful than assessing single or dual condition performance in isolation, as cost metrics incorporate individual differences in baseline performance (Verhaeghen et al.,2003).

Poorer dual task walking abilities have been related to increased fall risk (e.g., Bridenbaugh and Kressig,2015; Lundin-Olsson et al., 1997; Montero-Odasso et al.,2012), cognitive decline (Montero-Odasso et al.,2017), frailty, disability, and mortality (Verghese et al.,2012). Importantly, dual task walking performance is more predictive of falls in aging than single task walking performance (Ayers et al., 2014; Gillain et al., 2019; Halliday et al., 2018; Johansson et al., 2016; Verghese et al., 2017). This could be because dual task walking provides a better analog for real-world scenarios, such as talking to friends or reading street signs while walking. Indeed, a recent study reported that in-lab dual task walking attributes (gait speed, step regularity, and stride regularity) were more similar to real-world gait (measured during daily life with a wearable sensor), as compared with normal walking in lab with no dual tasking requirements (Hillel et al., 2019). Thus, given the link between dual task walking performance and falls, and its greater ecological validity, we selected to analyze dual instead of single task walking in the present work. There are clear cortical contributions to the control of walking (Allali et al., 2014; Koenraadt et al., 2014; Miyai et al., 2001; Petersen et al., 2012; Takakusaki, 2017). Thus, poorer dual task walking performance in older age has been attributed, at least in part, to age-related brain atrophy (Allali et al., 2019; Lucas et al., 2019; Ross et al., 2021). A large body of literature suggests that age-related structural brain atrophy occurs in an anterior-to-posterior pattern, with the frontal cortices atrophying earlier and faster than other regions of the brain (e.g., Fjell et al., 2009a; Lemaitre et al., 2012; Salat et al., 2004; Thambisetty et al., 2010). Given this, it is not surprising that previous work has linked lower prefrontal cortex gray matter volume with poorer dual task walking abilities in older adults (Tripathi et al., 2019; Wagshul et al., 2019). Aging is hypothesized to increase reliance on alternative (i.e., non-motor) neural resources, such as the frontal cortex (Mirelman et al., 2017), to compensate for brain atrophy in sensorimotor regions and maintain performance (Cabeza et al., 2002; Fettrow et al., 2021b; Steffener and Stern, 2012). Interestingly, recent work in a large sample of middle- to older-aged adults (*n* = 966) has reported disproportionately steep age differences (i.e., atrophy, demyelination, and iron reduction) in the sensorimotor cortices rather than in prefrontal regions (Taubert et al., 2020). Thus, structural changes in the sensorimotor cortices with aging may also contribute to age-related mobility declines.

Many previous studies have reported relationships between age differences in regional brain structure (e.g., atrophy in widespread cortical and subcortical regions, including the frontal and sensorimotor cortices, basal ganglia, cerebellum, and motor tracts) and worse gait for older adults during single task walking, such as slowed gait speed and increased gait variability (for review, see Tian et al., 2017; Wilson et al., 2019). However, compared to the extensive literature examining single task walking, only limited work examining brain structure has focused on dual task walking in aging. A majority of the studies examining correlates of dual task walking in aging have instead focused on brain function, using functional near-infrared spectroscopy (fNIRS). These studies have largely found increases in prefrontal cortex oxygenation levels from single to dual task walking for older adults, suggesting that dual compared with single task walking demands more prefrontal neural resources (e.g., Beurskens et al., 2014; Doi et al., 2013; Holtzer et al., 2015). As dual task walking is more cognitively demanding than normal walking, it is logical that functional contributions from the prefrontal cortex increase during dual task walking (Holtzer et al., 2015); thus, markers of prefrontal cortex structure might also relate to dual task walking performance in older age. Overall, while these functional studies provide important insight into the vasodynamic response to dual task walking, further work is needed to understand how markers of brain structure relate to dual task walking in aging as well.

The small body of work that has investigated relationships between brain structure and dual task walking in older adults suggests an important link between “maintenance” of brain structure and maintenance of dual task walking abilities. Two previous studies found associations between greater gait slowing during dual task walking in older adults and lower gray matter volume in the middle frontal gyrus (Allali et al., 2019), medial prefrontal and cingulate cortices, and thalamus (Tripathi et al., 2019). Further, several separate studies found that older adults who showed a greater increase in prefrontal cortex oxygenation from single to dual task walking also had lower white matter fractional anisotropy (averaged across the whole white matter mask; Lucas et al., 2019), lower gray matter volume within the frontal lobe (and specifically, the superior and rostral middle frontal gyri; Wagshul et al., 2019), and reduced cortical thickness across the frontal, parietal, temporal, occipital, cingulate, and insular cortices (Ross et al., 2021). These imaging metrics were not related to faster dual task walking, though, suggesting that the observed increases in prefrontal cortex activity represented compensation to maintain walking performance, despite atrophying brain structure.

The prior work described above examining the brain structural correlates of dual task walking tested only one structural imaging modality in isolation. Here we combined across multiple imaging modalities to provide more comprehensive information about age differences in brain structure and how these relate to dual task walking. We assessed volumetric metrics of atrophy, i.e., gray matter, cerebellar, hippocampal, and ventricular volume. We also examined surface metrics, including cortical thickness (Dahnke et al., 2013), sulcal depth (Yun et al., 2013), cortical complexity (i.e., folding complexity of the cortex; Yotter et al., 2011b), and gyrification index (i.e., mean curvature of the cortex; Luders et al., 2006). Surface-based morphometry metrics have several advantages over volume-based metrics (Hutton et al., 2009; Lemaitre et al., 2012; Winkler et al., 2010), including more accurate spatial registration (Desai et al., 2005), sensitivity to surface folding, and independence from head size (Gaser and Kurth, 2017). Despite these potential benefits, compared to volumetric measures, less work has examined how surface measures relate to dual task walking in aging.

We also examined white matter microstructure metrics derived from diffusion MRI, including free-water (FW) corrected fractional anisotropy (FAt, “t’ refers to the tissue compartment remaining after FW correction), axial diffusivity (ADt), and radial diffusivity (RDt), and the fractional volume of FW (Pasternak et al., 2009). FW correction is particularly important for analyses of older adult brains because age-related white matter degeneration can lead to enlarged interstitial spaces (Meier-Ruge et al., 1992) and thereby increased partial volume effects between white matter fibers and extracellular water (Chad et al., 2018). Recent work found that FW correction results in less pronounced age differences in white matter microstructure than previously reported (Chad et al., 2018), suggesting that prior age difference results are at least partially driven by fluid effects. Thus, to increase interpretability of white matter microstructural effects, it is important to correct for FW when examining white matter in aging. Moreover, higher FW has been related to poorer cognition in aging (Gullett et al., 2020; Maillard et al., 2019) and poorer function (e.g., bradykinesia) in Parkinson’s disease (Ofori et al., 2015).

In the present work, we addressed several aims: 1) To characterize age differences in brain structure; we predicted the most pronounced age differences in the prefrontal cortex. 2) To identify regions of age differences in the relationship between brain structure and DTcost of gait speed and variability; given the fNIRS literature reporting increased prefrontal cortex activation during dual task walking (Beurskens et al., 2014; Doi et al., 2013; Holtzer et al., 2015), we predicted that greater prefrontal atrophy would correlate with greater DTcost of gait speed and variability for older but not younger adults. 3) To determine the strongest predictors(s) of DTcost of gait in older adults using a stepwise regression approach. This was an exploratory aim, and thus we did not define an *a priori* hypothesis.

## 2 MATERIALS AND METHODS

The University of Florida’s Institutional Review Board provided ethical approval for the study. All individuals provided their written informed consent.

### 2.1 Participants

37 young and 25 older adults from the Gainesville, FL community participated in this study. Due to the coronavirus 2019 (COVID-19) global pandemic, data collection for this study was terminated early, before the planned sample size for older adult participants was attained. Two older adults were excluded from analyses of the *T*_1_-weighted images. One of these older adults did not fit within the 64-channel coil, so a 20-channel coil was used instead; due to low image quality, we excluded their data from further analysis. The other older adult *T*_1_-weighted scan was excluded due to an incidental brain tumor finding. Thus, *n* = 23 older adults for all analyses involving the *T*_1_-weighted images. Due to time constraints, a diffusion MRI was not collected for one young and two older adults; thus, *n* = 36 young and *n* = 21 older adults for all diffusion MRI analyses. Of note, we reported on a different subset of behavioral and brain metrics from this same cohort in two recent publications (Fettrow et al., 2021a; Hupfeld et al., 2021b).

We screened all subjects for MRI eligibility and, as part of the larger study, transcranial magnetic stimulation (TMS) eligibility. We excluded those with any MRI or TMS contraindications (e.g., implanted metal, claustrophobia, or pregnancy). We also excluded individuals with: history of any neurologic condition (e.g., stroke, Parkinson’s disease, seizures, or a concussion in the last six months); a current psychiatric condition (e.g., active depression or bipolar disorder); self-reported smokers; those who self-reported consuming more than two alcoholic drinks per day on average; and those with history of treatment for alcoholism. All participants were right-handed and self-reported their ability to walk unassisted for at least 10 minutes and to stand for at least 30 seconds with their eyes closed.

Prior to enrollment, we screened participants for suspected cognitive impairment over the phone using the Telephone Interview for Cognitive Status (TICS; de Jager et al., 2003). We excluded those who scored < 21 of 39 points; this is equivalent to scoring < 25 points on the Mini-Mental State Exam (MMSE) and indicates probable cognitive impairment (de Jager et al., 2003). At the first testing session, we re-screened participants for cognitive impairment using the Montreal Cognitive Assessment (MoCA; Nasreddine et al., 2005). We added one point to the scores of participants with ≤ 12 years of education (Nasreddine et al., 2005). We did not enroll those who scored < 23 of 30 points (Carson et al., 2018).

### 2.2 Testing Sessions

Before the first session, we collected self-reported participant information on: demographics (e.g., age, sex, and years of education), medical history, handedness, footedness, exercise, and sleep. We also collected anthropometric information (e.g., height, weight, and leg length). Participants then completed mobility testing, followed by an MRI scan approximately five days later (Fig. 1). For 24 hours prior to each session, participants were requested to not consume alcohol, nicotine, or any drugs other than the medications they disclosed to us. At the start of each session, participants completed the Stanford Sleepiness Questionnaire, which asks for self-report of the hours slept the previous night and a current sleepiness rating (Hoddes et al., 1972).

**Figure 1.**
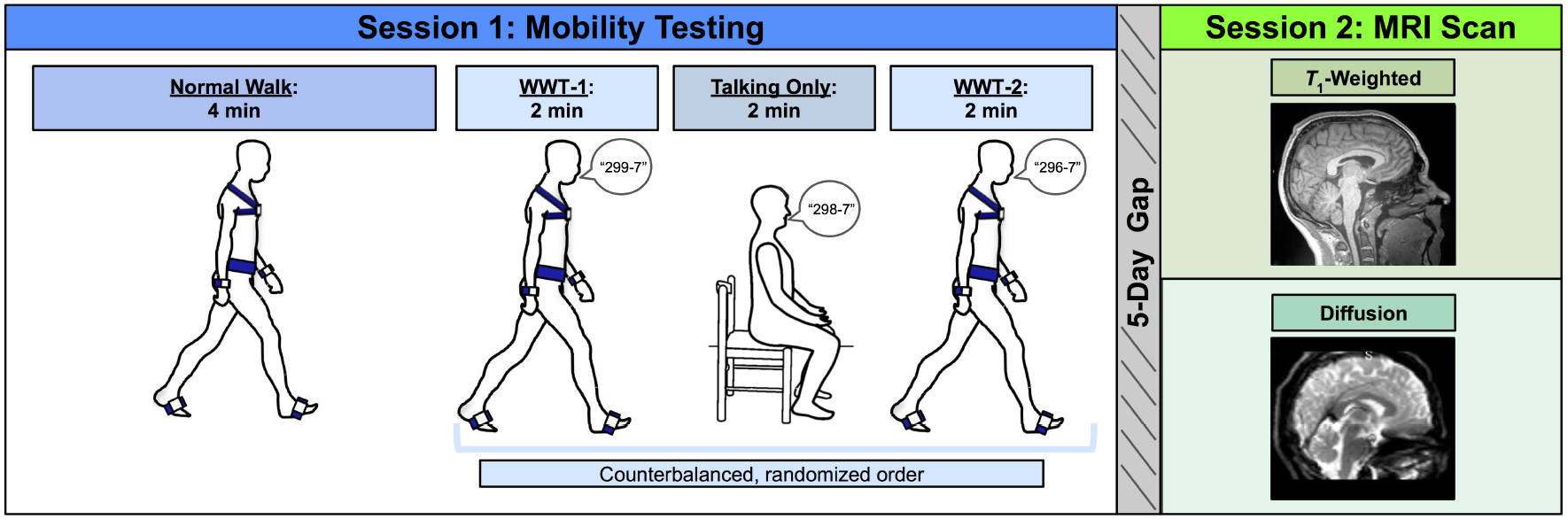
Methods overview. Left: During Session 1, participants first completed a normal (single task) overground walk (NW) at a comfortable self-selected speed. Next, participants completed three trials in a counterbalanced order: two walking while talking trials (WWT-1 and WWT-2) in which participants counted backwards by 7s while walking, and one talking only trial in which participants stayed seated while counting backwards by 7s. Right: Approximately five days later, during Session 2, participants completed an MRI protocol, which included a *T*_1_-weighted anatomical scan and a diffusion-weighted scan.

**Figure 2.**
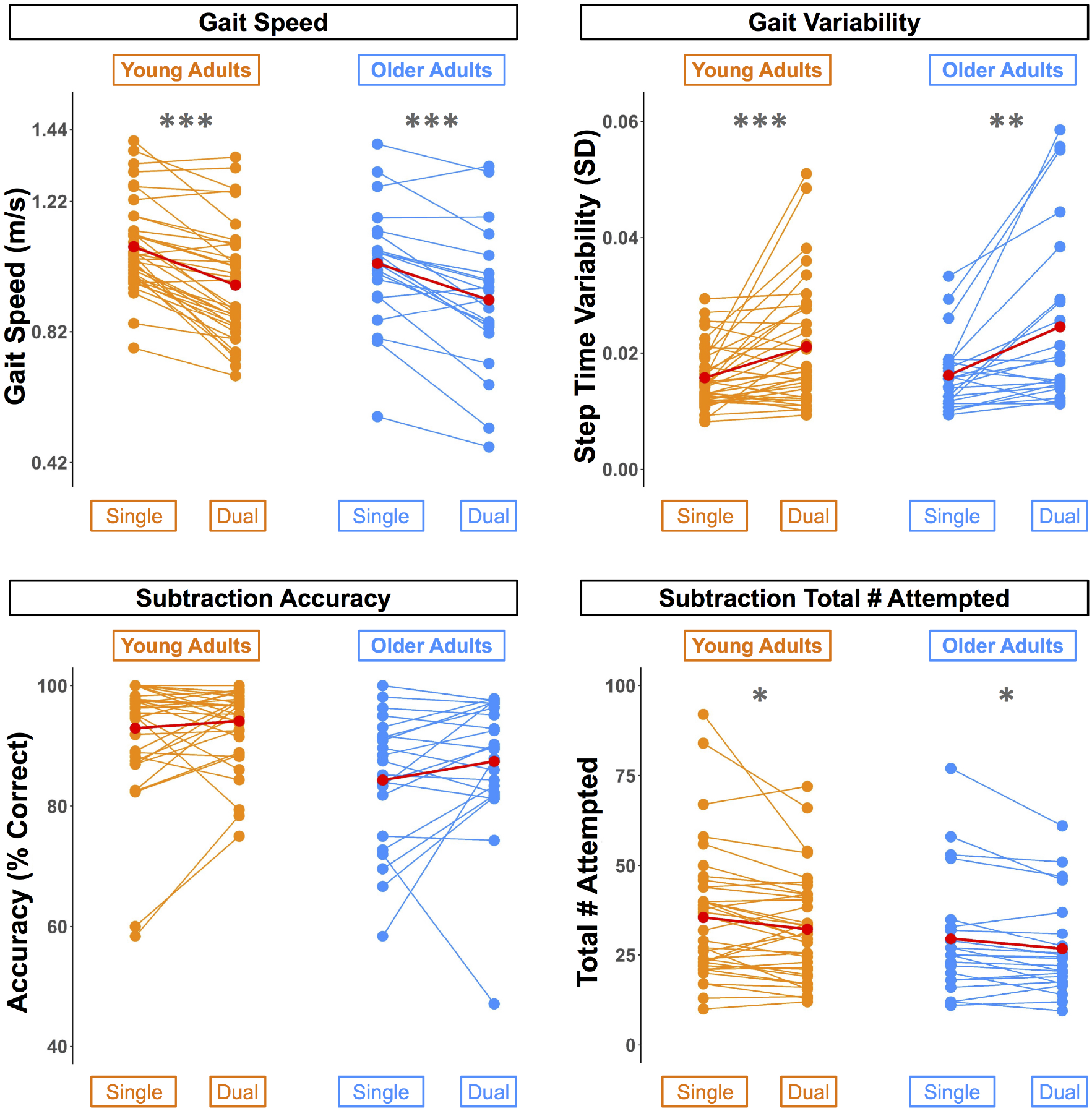
Differences in walking and subtraction performance during single versus dual task conditions. Gait and serial subtraction performance are depicted for each young (orange) and older (blue) adult. Each line represents one participant. Group means are shown in red. Across both age groups, gait speed slowed, gait variability increased, and number of subtraction problems attempted decreased from single to dual task conditions. \**p*_*FDR−corr*_ < 0.05, \*\**p*_*FDR−corr*_ < 0.01, \*\*\**p*_*FDR−corr*_ < 0.001.

### 2.3 Session 1: Mobility Testing

Participants completed three walking tasks while instrumented with six Opal inertial measurement units (IMUs; v2; APDM Wearable Technologies Inc., Portland, OR, USA). IMUs were placed on the feet, wrists, around the waist at the level of the lumbar spine, and across the torso at the level of the sternal angle (Fig. 1). First, participants walked back and forth across a 9.75 m room for four minutes at whichever pace they considered to be their “normal” walking speed (NW). Participants were instructed to refrain from talking, to keep their arms swinging freely at their sides, and to keep their head up and gaze straight ahead. Each time they reached the end of the room, they completed a 180-degree turn and walked the length of the room again.

Next, participants completed two trials of walking while talking (WWT-1 and WWT-2) and one trial of talking only. The WWT and talking only trials lasted for two minutes each. During the WWT trials, participants walked at their normal speed while counting backwards by 7s (Li et al., 2014), starting at number 299, 298, or 296. The WWT instructions were identical to those provided for the 4-minute walk, except that participants were additionally instructed to “try and pay equal attention to walking and talking” (Verghese et al., 2007). For the talking only trial, participants sat in a chair and counted backwards by 7s for two minutes. We counterbalanced the order of the WWT-1, WWT-2, and talking only trials and the starting number across all participants.

### 2.4 Spatiotemporal Variable Calculation

During both the walking and the balance tasks, we recorded inertial data using MobilityLab software (v2; APDM Wearable Technologies Inc., Portland, OR, USA). After each trial, MobilityLab calculated 14 spatiotemporal gait variables. The algorithm for calculating these metrics has been validated through comparison to force plate and motion capture data (see internal validation by MobilityLab: https://support.apdm.com/hc/en-us/articles/360000177066-How-are-Mobility-Lab-s-algorithms-validated- and (Washabaugh et al., 2017). To condense the gait variables into several summary metrics, for each trial, we extracted one variable from each of the four gait domains described by Hollman et al. (2011a): gait rhythm (cadence (steps/min)), gait phase (stance (% gait cycle)), gait pace (speed (m/s)), and gait variability (step time variability (standard deviation)). We calculated the average of each of these four variables for the NW and WWT-1 and WWT-2 trials to produce one variable for each of the four gait domains for NW and WWT.

### 2.5 Cognitive Outcome Variable Calculation

We also measured cognitive performance during the seated compared to WWT conditions. We examined both speed (i.e., total number of subtraction problems attempted) and accuracy (i.e., % correct) during both the seated and WWT conditions.

### 2.6 DTcost Calculation

To characterize differences in these gait and cognitive performance summary metrics between single and dual task conditions, similar to a large body of previous work (e.g., Kelly et al., 2010; Patel et al., 2014; Van Impe et al., 2011), we calculated the DTcost of each variable as follows:

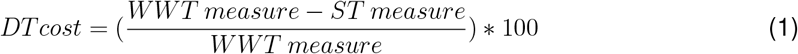

We then calculated a correlation matrix for the four resulting DTcost of gait measures across the whole sample. This revealed that DTcost of gait speed was highly correlated with the DTcost of cadence (r = 0.90, *p* < 0.001) and DTcost of stance time (r = −0.85, *p* < 0.001). Thus, we opted to analyze only two variables as primary outcome metrics in our final statistical analyses: 1) DTcost of gait speed; and 2) DTcost of step time variability. Both slower gait speed and increased step time variability have been related to higher fall risk for older adults (Callisaya et al., 2011; Espy et al., 2010; Quach et al., 2011).

### 2.7 Session 2: MRI Scan

We acquired an MRI scan for each participant using a Siemens MAGNETOM Prisma 3 T scanner (Siemens Healthcare, Erlangen, Germany) with a 64-channel head coil.

#### 2.7.1 Anatomical acquisition

We collected a 3D *T*_1_-weighted anatomical image using a magnetization-prepared rapid gradient-echo (MPRAGE) sequence. The parameters for this anatomical image were as follows: repetition time (TR) = 2000 ms, echo time (TE) = 3.06 ms, flip angle = 8°, field of view = 256 × 256 mm^2^, slice thickness = 0.8 mm, 208 slices, voxel size = 0.8 mm^3^.

#### 2.7.2 Diffusion-weighted acquisition

We also collected a diffusion-weighted spin-echo prepared echo-planar imaging sequence with the following parameters: 5 *b*_0_ scans (without diffusion weighting), 64 gradient directions with diffusion weighting 1000 s/mm^2^, TR = 6400 ms, TE = 58 ms, isotropic resolution = 2 × 2 × 2 mm, FOV = 256 × 256 mm^2^, 69 slices, phase encoding direction = Anterior to Posterior. Immediately prior to this acquisition, we collected 5 *b*_0_ scans (without diffusion weighting) in the opposite phase encoding direction (Posterior to Anterior) for later use in distortion correction.

### 2.8 *T*_1_-Weighted Image Processing for Voxelwise Analyses

#### 2.8.1 Gray matter volume

We processed the *T*_1_-weighted scans using the Computational Anatomy Toolbox toolbox (version r1725; Gaser et al., 2016; Gaser and Kurth, 2017) in MATLAB (R2019b). We implemented default CAT12 preprocessing steps, including the new adaptive probability region-growing skull stripping method. Briefly, the CAT12 pipeline includes segmentation into gray matter, white matter, and cerebrospinal fluid, followed by spatial normalization from subject space to standard space using high-dimensional Dartel registration and modulation. After CAT12 preprocessing was complete, we visually examined data quality by displaying each modulated, normalized gray matter segment and checking alignment between subjects and with the standard space template. We did not remove any scans as a result of visual inspection. All scans passed acceptable CAT12 quantitative quality control thresholds (i.e., resolution, noise, bias, and image quality > 80). Finally, we used the CAT12 *Check Sample Homogeneity* function to evaluate correlations between all gray matter segments. Gray matter segments for each participant were within two standard deviations of the group mean, indicating that the sample contained no outliers. To increase signal-to-noise ratio, we smoothed the modulated, normalized gray mattersegments using Statistical Parametric Mapping 12 (SPM12, v7771; Ashburner et al., 2014) with an 8 mm full width at half maximum kernel. We entered these preprocessed gray matter volume maps into the group-level voxelwise statistical models described in Section 2.13. We used CAT12 to calculate total intracranial volume for each participant for later use as a covariate in these group-level statistical analyses.

#### 2.8.2 Cortical surface metrics

The CAT12 pipeline also extracts surface-based morphometry metrics (Dahnke et al., 2013; Yotter et al., 2011a). To calculate surface metrics, CAT12 uses a projection-based thickness algorithm that handles partial volume information, sulcal blurring, and sulcal asymmetries without explicit sulcus reconstruction (Dahnke et al., 2013; Yotter et al., 2011a). We used CAT12 to extract four surface metrics: 1) cortical thickness: the thickness of the cortical gray matter between the outer surface (i.e., the gray matter-cerebrospinal fluid boundary) and the inner surface (i.e., the gray matter-white matter boundary) (Dahnke et al., 2013); 2) cortical complexity: fractal dimension, a metric of folding complexity of the cortex (Yotter et al., 2011b); 3) sulcal depth: the Euclidean distance between the central surface and its convex hull (Yun et al., 2013); and 4) gyrification index: a metric based on the absolute mean curvature, which quantifies the amount of cortex buried within the sulcal folds as opposed to the amount of cortex on the “outer” visible surface (Luders et al., 2006). Prior to further analysis, we visually checked all cortical surface data using CAT12’s *Display Surfaces* tool and then resampled and smoothed the surfaces at 15 mm for cortical thickness and 20 mm for the three other metrics. We entered these resampled and smoothed surface files into the group-level voxelwise statistical models described in Section 2.13.

#### 2.8.3 Cerebellar volume

To improve the normalization of the cerebellum (Diedrichsen, 2006; Diedrichsen et al., 2009), similar to our past work (Hupfeld et al., 2021a; Salazar et al., 2020, 2021), we applied specialized preprocessing steps to the cerebellum to produce cerebellar volume maps. First, we entered each participant’s whole-brain *T*_1_-weighted image into the CEREbellum Segmentation (CERES) pipeline (Romero et al., 2017). CERES uses a patch-based segmentation approach to segment the cerebellum from the cortex; this automated method has been demonstrated to perform better than either semi-automatic or manual cerebellar segmentation (Romero et al., 2017). We visually inspected the resulting segmentations, created a binary mask from each participant’s CERES cerebellar segmentation, and used this mask to extract their cerebellum from their whole-brain *T*_1_-weighted image. We then used rigid, affine, and Symmetric Normalization (SyN) transformation procedures within the Advanced Normalization Tools package (ANTs; v1.9.17; Avants et al., 2010, 2011) to warp (in a single step) each participant’s extracted subject space cerebellum to a 1 mm cerebellar template in standard space, the Spatially Unbiased Infratentorial Template (SUIT) template (Diedrichsen, 2006; Diedrichsen et al., 2009). The SUIT template was selected because it offers greater detail of internal cerebellar structures compared to whole brain templates, which improves cerebellar normalization (Diedrichsen, 2006; Diedrichsen et al., 2009). For this warping we used a version of the SUIT template with the brainstem removed, as the CERES cerebellar segmentation does not include the brainstem.

The flowfields that were applied to warp these cerebellar segments to SUIT space were additionally used to calculate the Jacobian determinant image, using ANTs’ *CreateJacobianDeterminantImage*.*sh* function; the Jacobian determinant encodes local shrinkage and expansion for each voxel between subject space and the target image (i.e., here, the standard space template). We multiplied each normalized cerebellar segment by its corresponding Jacobian determinant to produce modulated cerebellar images in standard space for each participant. Modulation preserves the volumes present in the original untransformed (subject space) image. Lastly, to increase signal-to-noise ratio, we smoothed the modulated, normalized cerebellar images using a kernel of 2 mm full width at half maximum and entered the resulting cerebellar volume maps into the group-level voxelwise statistical models described in Section 2.13. Of note, we used cerebellar total volumes in our analyses instead of segmenting the cerebellum by tissue type, in order to avoid any inaccuracy due to low contrast differences between cerebellar gray and white matter.

### 2.9 Diffusion-Weighted Image Processing for Voxelwise Analyses

#### 2.9.1 Diffusion preprocessing

See Supplemental Information for further details regarding preprocessing of the diffusion-weighted data. We first visually inspected raw scans for artifacts and excessive head movement. We then corrected images for signal drift (Vos et al., 2017) using the ExploreDTI graphical toolbox (v4.8.6; www.exploredti.com; Leemans et al., 2009) in MATLAB (R2019b). Next, we used the FMRIB Software Library (FSL; v6.0.1; Jenkinson et al., 2012; Smith et al., 2004) processing tool *topup* to estimate the susceptibility-induced off-resonance field (Andersson et al., 2003). This procedure yielded a single corrected field map for use in eddy current correction. We used FSL’s *eddy _ cuda* to simultaneously correct the data for eddy current-induced distortions and both inter- and intra-volume head movement (Andersson and Sotiropoulos, 2016).

#### 2.9.2 FW correction and tensor fitting

We implemented a custom FW imaging algorithm (Pasternak et al., 2009) in MATLAB. This algorithm estimates FW fractional volume and FW corrected diffusivities by fitting a two-compartment model at each voxel (Pasternak et al., 2009). The two-compartment model consists of: 1) a tissue compartment modeling water molecules within or in the vicinity of white matter tissue, quantified by diffusivity (FAt, RDt, and ADt); and 2) a FW compartment, reflecting the proportion of water molecules with unrestricted diffusion, and quantified by the fractional volume of this compartment. FW ranges from 0 to 1; FW = 1 indicates that a voxel is filled with freely diffusing water molecules (e.g., as in the ventricles). These metrics (FAt, RDt, ADt, FW) are provided as maps for each voxel in the brain.

### 2.10 Tract-Based Spatial Statistics

We applied FSL’s tract-based spatial statistics (TBSS) processing steps to prepare the data for voxelwise analyses across participants (Smith et al., 2006). Benefits of TBSS include avoiding problems associated with suboptimal image registration between participants and eliminating the need for spatial smoothing. TBSS uses a carefully-tuned nonlinear registration and projection onto an alignment-invariant tract representation (i.e., the mean FA skeleton); this process improves the sensitivity, objectivity, and interpretability of analyses of multi-subject diffusion studies. We used the TBSS pipeline as provided in FSL, which first includes eroding the FA images slightly and zeroing the end slices. Next, each participant’s FA data is brought into a common space (i.e., the FMRIB58 FA 1 mm isotropic template) using the nonlinear registration tool FNIRT (Andersson et al., 2007b,a). A mean FA image is then calculated and thinned to create a mean FA skeleton. Then, each participant’s aligned FA data is projected onto the group mean skeleton. Lastly, we applied the same nonlinear registration to the FW, FAt, RDt, and ADt maps to project these data onto the original mean FA skeleton. Ultimately, these TBSS procedures resulted in skeletonized FW, FAt, ADt, and RDt maps in standard space for each participant. These were the maps that we entered in the group-level voxelwise statistical models described in Section 2.13.

### 2.11 Image Processing for Region of Interest Analyses

#### 2.11.1 Ventricle and gray matter volume regions of interest

CAT12 automatically calculates the inverse warp, from standard space to subject space, for several volume-based atlases. We isolated multiple regions of interest (ROIs) from these atlases in subject space: the lateral ventricles and pre- and postcentral gyri from the Neuromorphometrics (http://Neuromorphometrics.com) volume-based atlas, and the thalamus, striatum, and globus pallidus from the CoBra Subcortical atlas (Tullo et al., 2018; Fig. S1). We visually inspected each ROI mask overlaid onto each participant’s *T*_1_-weighted image in ITK-SNAP and hand corrected the ROI mask if needed (Yushkevich et al., 2006). Using *fslstats*, we extracted the number of voxels in each ROI mask in subject space and calculated the mean image intensity within the ROI in the subject space cerebrospinal fluid (lateral ventricles) or gray matter segment (for all of the other ROIs). We then calculated ROI volume in mL as: (number of voxels in the ROI mask)*(mean intensity of the tissue segment within the ROI mask)*(volume/voxel). In subsequent statistical analyses, we used the average of the left and right side structures for each ROI, and we entered these ROI volumes as a percentage of total intracranial volume (to account for differences in head size).

#### 2.11.2 FW ROIs

We also extracted FW values from the diffusion MRI maps for the same ROIs for which we calculated gray matter volume. We rigidly registered the subject space *T*_1_-weighted image to the subject space FW image. (We used a rigid registration in this case because we previously used *topup* to resolve distortions during DWI preprocessing; Section 2.9.1). We then used ANTs to apply the inverse of that transformation to the subject *T*_1_-space atlases described in Section 2.11.1. This resulted in volumetric atlases for each participant in their native diffusion space. We then isolated masks for the same ROIs described in Section 2.11.1 from these atlases and visually inspected each ROI mask overlaid onto each participant’s FW map in ITK-SNAP. Finally, we used *fslstats* to extract mean image intensity in the FW map within each ROI mask. Here we used mean intensity as our outcome metric (rather than volume in mL as above) to estimate the fractional volume of FW within the ROI and obtain a metric more representative of microstructural FW, rather than the size of the ROI which represents macrostructural atrophy. We calculated the average mean intensity for the left and right side for each structure and used this average value in subsequent statistical analyses.

#### 2.11.3 Hippocampal ROIs

We implemented the Automatic Segmentation of Hippocampal Subfields (ASHS)-T1 (Yushkevich et al., 2015) pipeline within ITK-SNAP (Yushkevich et al., 2015) to segment and extract the volume in mL of three hippocampal structures: anterior hippocampus, posterior hippocampus, and parahippocampal cortex. The ASHS pipeline uses a multi-atlas segmentation framework and super-resolution approach; this outperforms alternative *T*_1_ hippocampal segmentation pipelines by reducing misclassification of meninges as gray matter(Yushkevich et al., 2015). Though this pipeline is currently validated for use on only older adults (defined as those 55+ years old; Yushkevich et al., 2015), for completeness, here we also implemented the pipeline onmyyounger adult participants. For statistical analyses, we used the average of the left and right side structures, and we entered these volumes as a percentage of total intracranial volume (to account for differences in head size).

### 2.12 Statistical Analyses

#### 2.12.1 Participant characteristics, testing timeline, and mobility performance

We conducted all statistical analyses on the demographic and behavioral data using using R (v4.0.0; R Core Team, 2013). For each set of analyses, we applied the Benjamini-Hochberg false discovery rate (FDR) correction to the *p* values for the age group predictor (Benjamini and Hochberg, 1995).

#### 2.12.2 Demographic and behavioral data

First, we compared demographic, physical characteristics, and testing timeline variables between the age groups. We tested the parametric t-test assumptions: normality within each group (Shapiro test, *p* > 0.05) and homogeneity of variances between groups (Levene’s test, *p* > 0.05). The majority of variables did not meet parametric assumptions, so we conducted nonparametric two-sided Wilcoxon rank-sum tests for age group differences. We report the group medians and interquartile ranges for each of these variables. We also report nonparametric effect sizes (Field et al., 2012; Rosenthal et al., 1994). To test for differences in the sex distribution within each age group, we conducted a Pearson chi-square test.

#### 2.12.3 Age differences in the DTcost of gait and subtraction performance

To examine whether gait and subtraction performance differed between the single and dual task conditions and/or between the age groups, we used a linear mixed model approach (lme; Pinheiro et al., 2007). We entered age group, condition (i.e., single or dual task), and the age group*condition interaction as predictors, and included a random intercept for each subject.

### 2.13 Voxelwise Statistical Models

We tested the same voxelwise models for each of the imaging modalities. In each case, we defined the model using SPM12 and then re-estimated each model using the Threshold-Free Cluster Enhancement toolbox (TFCE; http://dbm.neuro.uni-jena.de/tfce) with 5,000 permutations. This toolbox provides non-parametric estimation using TFCE for models previously estimated using SPM parametric designs. Statistical significance was determined at *p* < 0.05, family-wise error (FWE) corrected for multiple comparisons.

#### 2.13.1 Age differences

First, we conducted two-sample t-tests to test for age differences in brain structure. In each of these models, we set the imaging modality (e.g., normalized, modulated gray matter volume segments) as the outcome variable and controlled for sex. In the gray matter and cerebellar volume models, we also controlled for head size (i.e., total intracranial volume). Also in the gray matter volume models only, we set the absolute masking threshold to 0.1 (Gaser and Kurth, 2017) and used an explicit gray matter mask that excluded the cerebellum (because we analyzed cerebellar volume separately from “whole brain” gray matter volume; Section 2.8.3).

#### 2.13.2 Interaction of age group * DTcost of gait

Our primary analysis of interest tested for regions in which the relationship between brain structure and the DTcost of gait differed between young and older adults. We ran two-group t-test models and included the DTcost of gait speed or step time variability for young and older adults as covariates of interest. We tested for regions in which the correlation between brain structure and DTcost was greater for the young compared with the older adults, and where the correlation between brain structure and DTcost was lower for the young compared with the older adults. As above, we controlled for sex in all models, and we controlled for head size in the gray matter and cerebellar volume models.

### 2.14 ROI Statistical Models

We conducted ROI analyses in R. For each set of analyses, we applied the Benjamini-Hochberg FDR correction to the *p* values for the predictor(s) of interest (Benjamini and Hochberg, 1995).

#### 2.14.1 Age differences

Similar to the above voxelwise models, we first ran linear models to test for age group differences in ROI volume or mean intensity, controlling for sex. We applied the FDR correction to the *p* values for the age group predictor (i.e., the primary analysis of interest). *Post hoc*, we also FDR-corrected the *p* values for the sex predictor, to better interpret several statistically significant sex difference results.

#### 2.14.2 Interaction of age group * DTcost of gait

Also similar to above, we ran linear models testing for an interaction of age group with the DTcost of gait speed or step time variability, controlling for sex. We FDR-corrected the *p* values for the interaction term.

### 2.15 Multiple Regression to Identify the Best Predictors of DTcost of Gait in Older Adults

We used two stepwise multivariate linear regressions to directly compare the neural correlates of the DTcost of gait identified by the voxelwise and ROI analyses described above. We ran one model for the DTcost of gait speed, and one model for the DTcost of step time variability. We included only the older adults in these models because the older adults showed stronger relationships between brain structure and the DTcost of gait (whereas the young adults tended to show either a weak relationship or no clear relationship between brain structure and the DTcost of gait).

In each of the two full models, we included sex and values from the peak result coordinate for each voxelwise model that indicated a statistically significant age difference in the relationship between brain structure and the DTcost of gait as predictors. We also included ROI values as predictors in any cases where the linear model yielded a significant age group by DTcost interaction term. We used *stepAIC* (Venables et al., 1999) *to produce a final model that retained only the best predictor variables; stepAIC* selects a maximal model based on the combination of predictors that produces the smallest Akaike information criterion (AIC). Overall, this stepwise regression approach allowed us to fit the best models using brain structure to predict the DTcost of gait for the older adults.

### 2.16 Comparison of Participant Characteristics and Testing Timeline

There were no statistically significant differences between the age groups in sex, handedness, footedness, alcohol use, or hours of sleep prior to each testing session. There were also no age group differences in the number of days elapsed between the testing sessions or in the difference in start time for the sessions. Older adults did report higher body mass indices, less physical activity, lower balance confidence, and greater fear of falling compared with young adults. See Table 1 for complete demographic information.

**Table 1.**
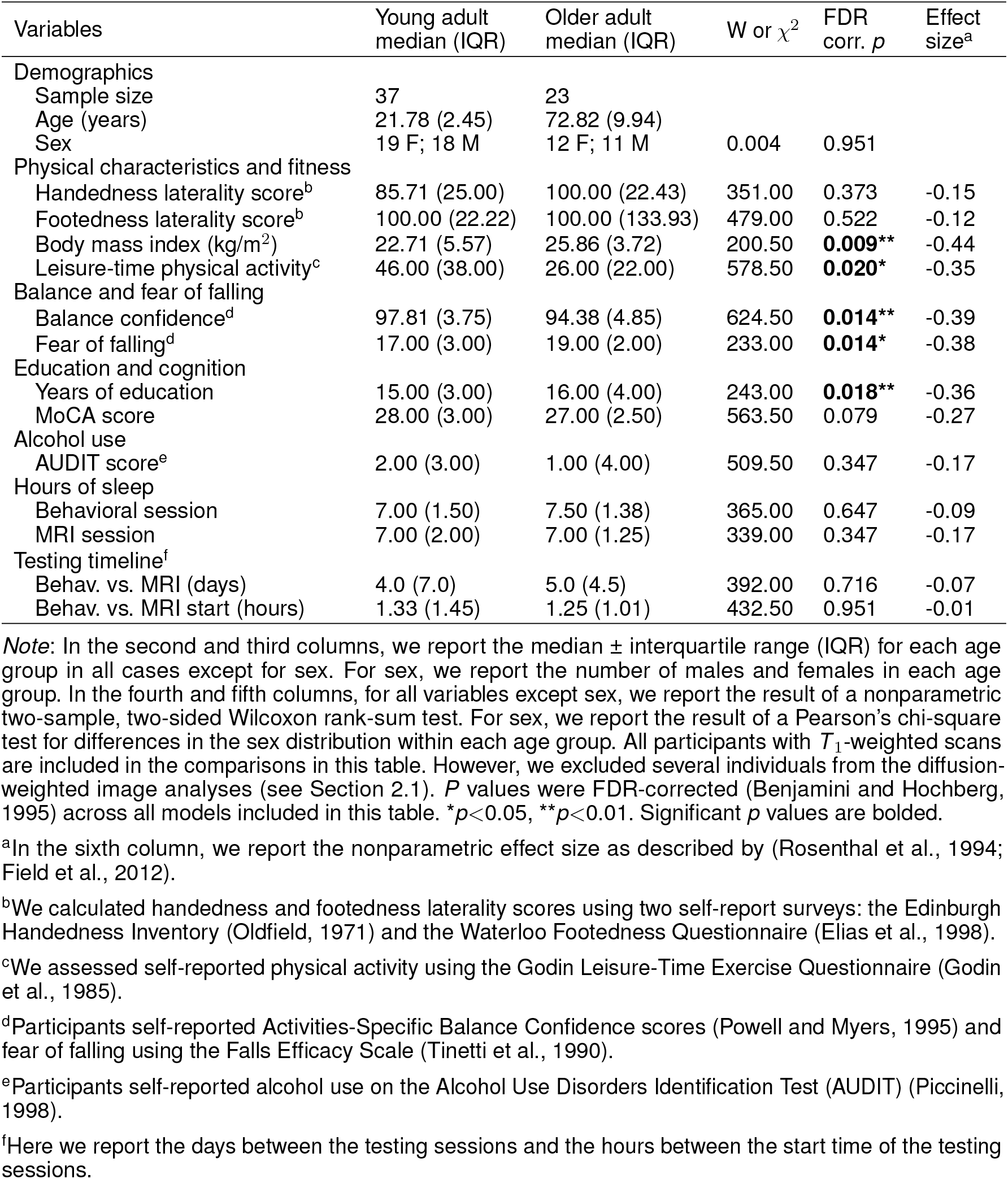
Participant characteristics and testing timeline

| Variables | Young adult<br>median (IQR) | Older adult<br>median (IQR) | W or $\chi^2$ | FDR<br>corr. $p$ | Effect<br>size<sup>a</sup> |
| --- | --- | --- | --- | --- | --- |
| Demographics |  |  |  |  |  |
| Sample size | 37 | 23 |  |  |  |
| Age (years) | 21.78 (2.45) | 72.82 (9.94) |  |  |  |
| Sex | 19 F; 18 M | 12 F; 11 M | 0.004 | 0.951 |  |
| Physical characteristics and fitness |  |  |  |  |  |
| Handedness laterality score<sup>b</sup> | 85.71 (25.00) | 100.00 (22.43) | 351.00 | 0.373 | -0.15 |
| Footedness laterality score<sup>b</sup> | 100.00 (22.22) | 100.00 (133.93) | 479.00 | 0.522 | -0.12 |
| Body mass index (kg/m<sup>2</sup>) | 22.71 (5.57) | 25.86 (3.72) | 200.50 | **0.009\*\*** | -0.44 |
| Leisure-time physical activity<sup>c</sup> | 46.00 (38.00) | 26.00 (22.00) | 578.50 | **0.020\*** | -0.35 |
| Balance and fear of falling |  |  |  |  |  |
| Balance confidence<sup>d</sup> | 97.81 (3.75) | 94.38 (4.85) | 624.50 | **0.014\*\*** | -0.39 |
| Fear of falling<sup>d</sup> | 17.00 (3.00) | 19.00 (2.00) | 233.00 | **0.014\*** | -0.38 |
| Education and cognition |  |  |  |  |  |
| Years of education | 15.00 (3.00) | 16.00 (4.00) | 243.00 | **0.018\*\*** | -0.36 |
| MoCA score | 28.00 (3.00) | 27.00 (2.50) | 563.50 | 0.079 | -0.27 |
| Alcohol use |  |  |  |  |  |
| AUDIT score<sup>e</sup> | 2.00 (3.00) | 1.00 (4.00) | 509.50 | 0.347 | -0.17 |
| Hours of sleep |  |  |  |  |  |
| Behavioral session | 7.00 (1.50) | 7.50 (1.38) | 365.00 | 0.647 | -0.09 |
| MRI session | 7.00 (2.00) | 7.00 (1.25) | 339.00 | 0.347 | -0.17 |
| Testing timeline<sup>f</sup> |  |  |  |  |  |
| Behav. vs. MRI (days) | 4.0 (7.0) | 5.0 (4.5) | 392.00 | 0.716 | -0.07 |
| Behav. vs. MRI start (hours) | 1.33 (1.45) | 1.25 (1.01) | 432.50 | 0.951 | -0.01 |

*Note:* In the second and third columns, we report the median ± interquartile range (IQR) for each age group in all cases except for sex. For sex, we report the number of males and females in each age group. In the fourth and fifth columns, for all variables except sex, we report the result of a nonparametric two-sample, two-sided Wilcoxon rank-sum test. For sex, we report the result of a Pearson's chi-square test for differences in the sex distribution within each age group. All participants with $T_1$ -weighted scans are included in the comparisons in this table. However, we excluded several individuals from the diffusion-weighted image analyses (see Section 2.1). $P$ values were FDR-corrected (Benjamini and Hochberg, 1995) across all models included in this table. \* $p < 0.05$ , \*\* $p < 0.01$ . Significant $p$ values are bolded.
<sup>a</sup>In the sixth column, we report the nonparametric effect size as described by (Rosenthal et al., 1994; Field et al., 2012).
<sup>b</sup>We calculated handedness and footedness laterality scores using two self-report surveys: the Edinburgh Handedness Inventory (Oldfield, 1971) and the Waterloo Footedness Questionnaire (Elias et al., 1998).
<sup>c</sup>We assessed self-reported physical activity using the Godin Leisure-Time Exercise Questionnaire (Godin et al., 1985).
<sup>d</sup>Participants self-reported Activities-Specific Balance Confidence scores (Powell and Myers, 1995) and fear of falling using the Falls Efficacy Scale (Tinetti et al., 1990).
<sup>e</sup>Participants self-reported alcohol use on the Alcohol Use Disorders Identification Test (AUDIT) (Piccinelli, 1998).
<sup>f</sup>Here we report the days between the testing sessions and the hours between the start time of the testing sessions.

### 2.17 Age and Condition Differences in Performance

Across both age groups, gait speed slowed and gait variability increased during WWT compared to NW (Table 2; Fig. S2). There was not a statistically significant difference in serial subtraction accuracy between the seated and WWT conditions (Table 2), though both young and older adults attempted fewer subtraction problems during the WWT conditions compared to the seated condition (Table 2; Fig. S2). Thus, across both age groups, subtraction speed decreased from single to dual task, but accuracy did not change.

**Table 2.**
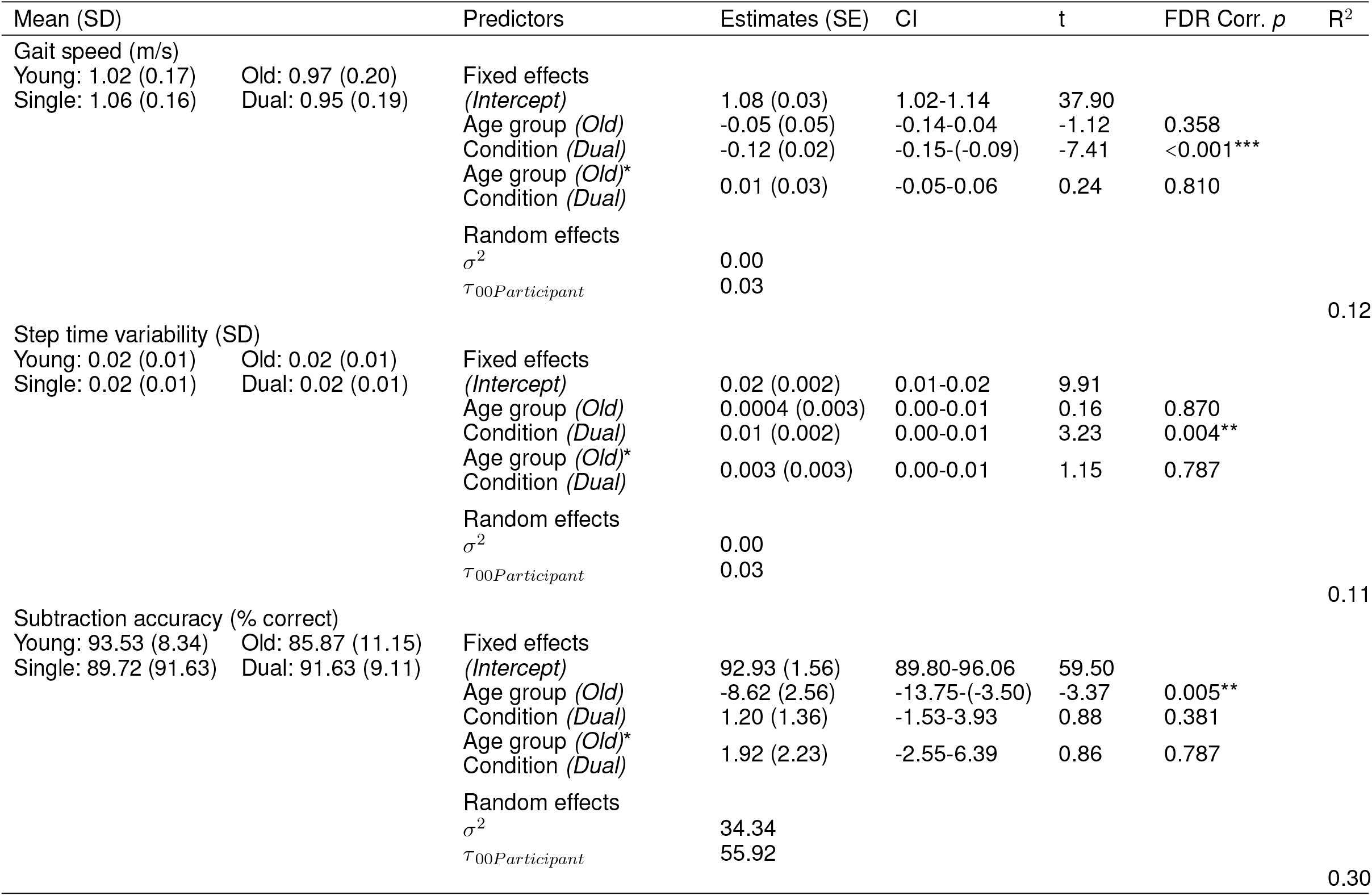
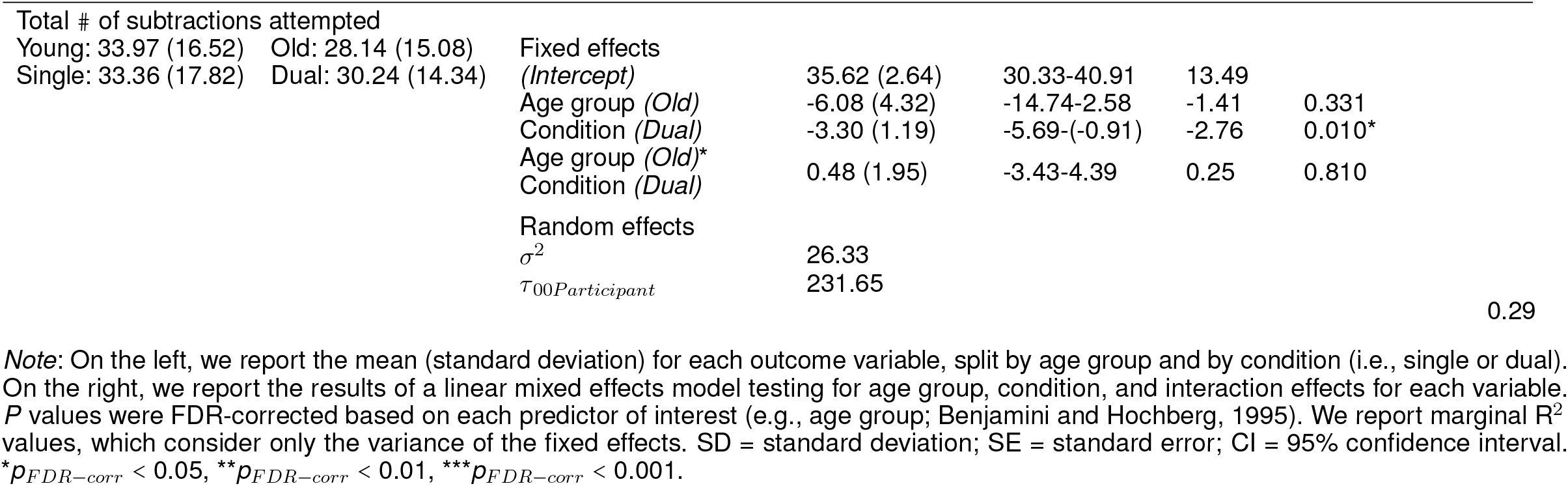
Age and condition differences in gait and subtraction performance

Across both conditions, the young adults performed with higher accuracy compared with the older adults (Table 2). However, there were no statistically significant age group differences in the DTcost of walking or subtraction performance (i.e., there were no significant age group by condition interactions; 2; Fig. S2). That is, the magnitude of single to dual task decrements in gait speed and number of subtraction problems attempted, as well as the magnitude of the increase in gait variability, was similar for young and older adults.

### 2.18 Comparison of Brain Structure Between Age Groups

#### 2.18.1 *T*_1_-weighted MRI metrics

Across the whole brain, older adults had significantly lower gray matter volume compared with young adults (Fig. 3). The greatest differences between young and older adults occurred in the bilateral pre- and postcentral gyri, temporal lobe, insula, and inferior portion of the frontal cortex. Cerebellar volume was lower for older compared with younger adults across most of the cerebellum, though there were no age differences in some regions, including the vermis and bilateral crus I (Fig. 3). Across the entire cortical surface, older adults had lower cortical thickness compared with young adults (Fig. 4). The largest age differences in cortical thickness occurred in the bilateral pre- and postcentral gyri and portions of the superior frontal cortex. Gyrification index was lower for older adults in the bilateral insula only. Cortical complexity was lower for older adults across portions of the bilateral insula, left middle frontal cortex, and posterior cingulate gyrus. Sulcal depth was reduced for older adults across the bilateral temporal lobes and insula, within the lateral fissure of the brain. Sulcal depth was higher for older compared with young adults across the superior frontal cortex, along the midline (Fig. 4).

**Figure 3.**
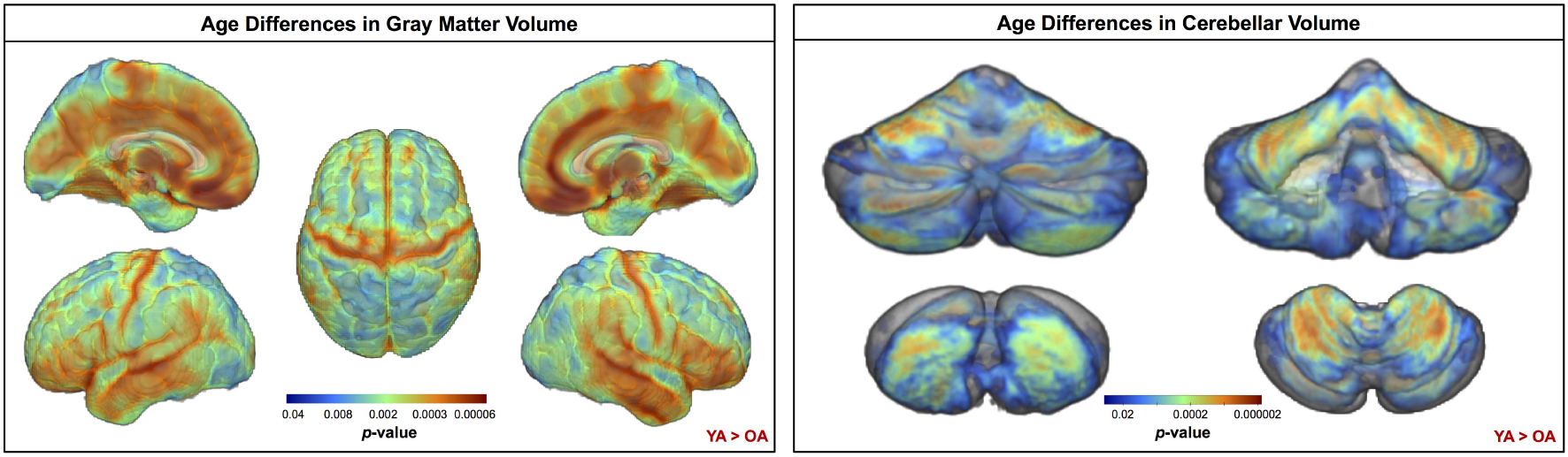
Age differences in gray matter and cerebellar volume. Increasingly warm colors indicate regions where young adult volumes were greater than older adult volumes. Results are overlaid onto a whole brain MNI-space template (left) and onto the SUIT cerebellar template (right). *p*_*FWE−corr*_ < 0.05.

**Figure 4.**
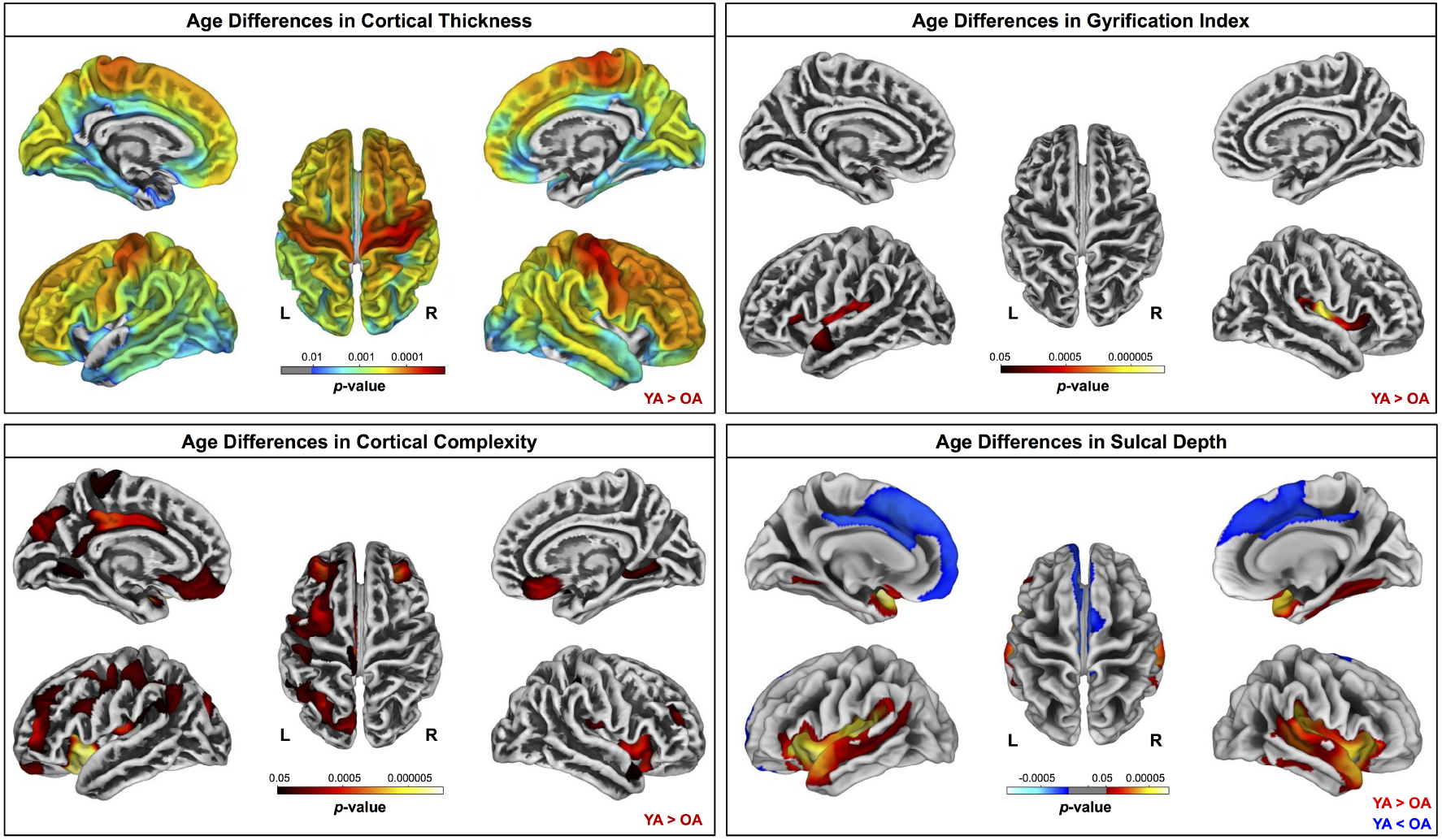
Age differences in surface measures. Warm colors indicate regions where young adult values were greater than older adult values. Cool colors indicate regions where young adult values were lower than older adult values. Results are overlaid onto CAT12 standard space templates. L = left; R = right. *p*_*FWE−corr*_ < 0.05.

### 2.19 Diffusion MRI Metrics

Compared with young adults, older adults showed lower FAt, lower ADt, higher RDt, and higher FW across almost the entire white matter skeleton (Fig. 5). There were some exceptions to this pattern, however, in portions of the superior corona radiata, corpus callosum (e.g., splenium), internal capsule, and thalamic radiations in which older adults showed higher FAt, higher ADt, and lower RDt compared with young adults.

**Figure 5.**
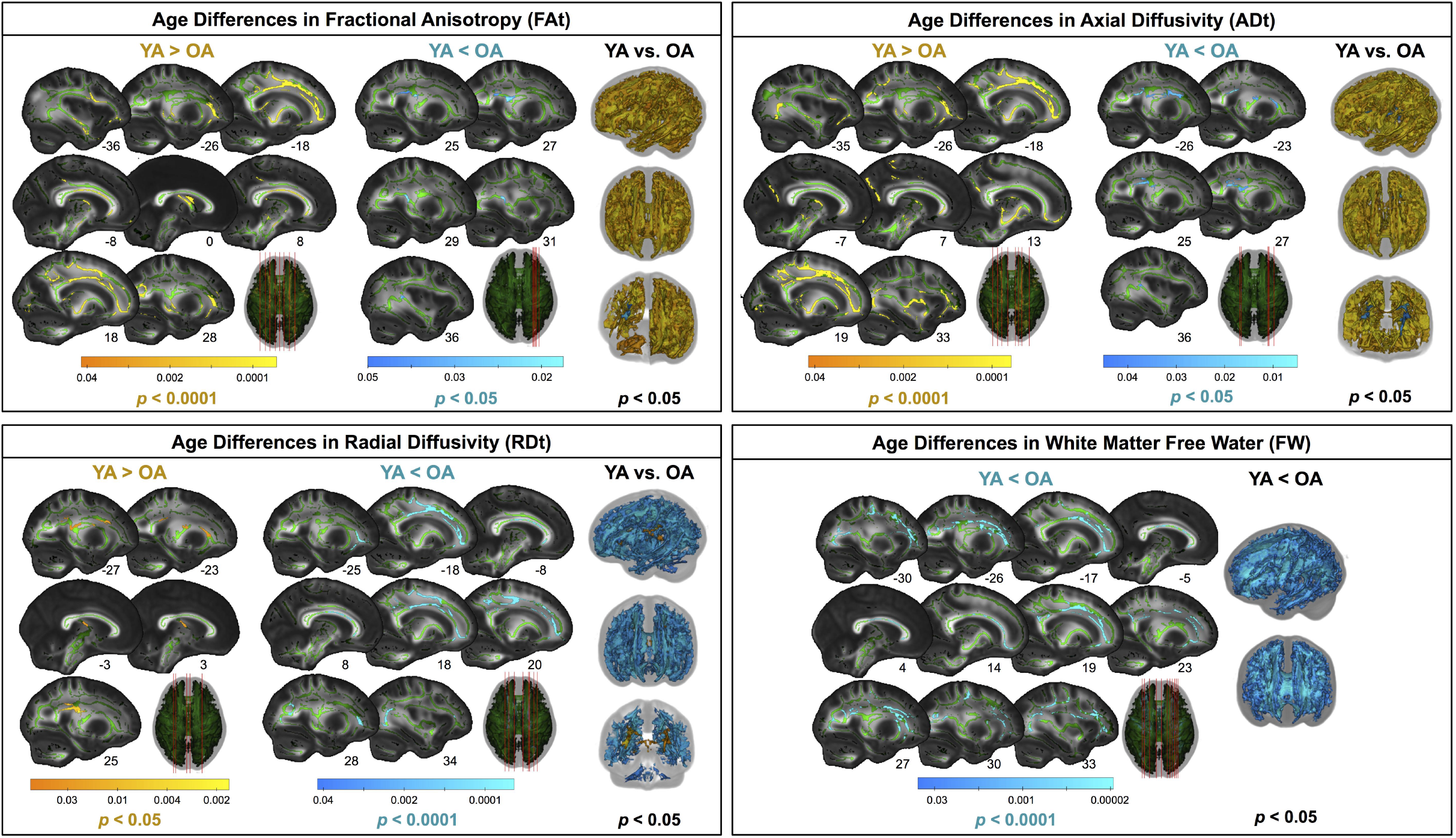
Age differences in FW-corrected white matter microstructure. Warm colors indicate regions where young adult values were greater than older adult values. Cool colors indicate regions where young adult values were lower than older adult values. Results are shown on the FMRIB58 FA template with the group mean white matter skeleton (green) overlaid. Age differences at *p*_*FWE−corr*_ < 0.05 covered almost the entire white matter skeleton; these results are depicted in the rightmost column of each panel. The left portion of each panel depicts more conservative statistical thresholding (noted under each colorbar) to better illustrate which regions showed the most pronounced age differences.

### 2.20 ROIs

Lateral ventricular volume was higher for older compared with younger adults (Table S1; Fig. S3). Older adults exhibited lower gray matter volume in all ROIs except for the globus pallidus and higher FW in all ROIs except for postcentral gyrus (Table S1; Fig. S4). Older adults had lower hippocampal volume across each of the three hippocampal ROIs (Table S1; Fig. S5). In several regions, pooling across both age groups, females had higher gray matter volume (thalamus) and FW (pre- and postcentral gyri and thalamus) compared with males.

### 2.21 Age Differences in the Relationship of Brain Structure with the DTcost of Gait Speed

There were no statistically significant age group by DTcost of gait speed interactions for gray matter or cerebellar volume. However, for the older adults, shallower sulcal depth across the sensorimotor, supramarginal, and superior frontal and parietal cortices was associated with greater DTcost of gait speed (Fig. 6; Table 3). That is, those older adults who showed the largest decreases in gait speed from single to dual task also had the shallowest sulcal depth across these regions. Young adults did not exhibit a clear relationship between sulcal depth in these regions and the DTcost of gait speed. There were no statistically significant age group differences in the correlation of cortical thickness, cortical complexity, or gyrification index with the DTcost of gait speed.

**Table 3.** Regions of age difference in the relationship of sulcal depth with the DTcost of gait speed and step time variability

| Region | Overlap of Atlas Region | TFCE Level |  |
| --- | --- | --- | --- |
| | | Extent ($k_E$) | $p_{FWE-corr}$ |
| DTcost of gait speed |  |  |  |
| L precentral gyrus | 31% | 3573 | 0.012* |
| L postcentral gyrus | 25% | — | — |
| L supramarginal gyrus | 19% | — | — |
| L superior frontal gyrus | 15% | — | — |
| L superior parietal lobule | 100% | 196 | 0.048* |
| DTcost of step time variability |  |  |  |
| L precentral gyrus | 25% | 5720 | 0.008** |
| L postcentral gyrus | 20% | — | — |
| L supramarginal gyrus | 17% | — | — |
| L insula | 8% | — | — |
| L pars opercularis | 7% | — | — |
| L pars triangularis | 6% | — | — |
| L superior parietal lobule | 5% | — | — |
| L superior frontal gyrus | 5% | — | — |

*Note:* Here we list all atlas regions from the Desikan-Killiany DK40 atlas (Desikan et al., 2006) that overlapped by 5% or more with each resulting cluster. The clusters were sorted by $p_{FWE-corr}$ value (from smallest to largest), then by cluster size (from largest to smallest). We do not list volumetric (e.g., MNI space) coordinates in this table because volumetric coordinates cannot be mapped directly onto cortical surfaces. L = left. \* $p_{FWE-corr} < 0.05$ , \*\* $p_{FWE-corr} < 0.01$ .

**Figure 6.**
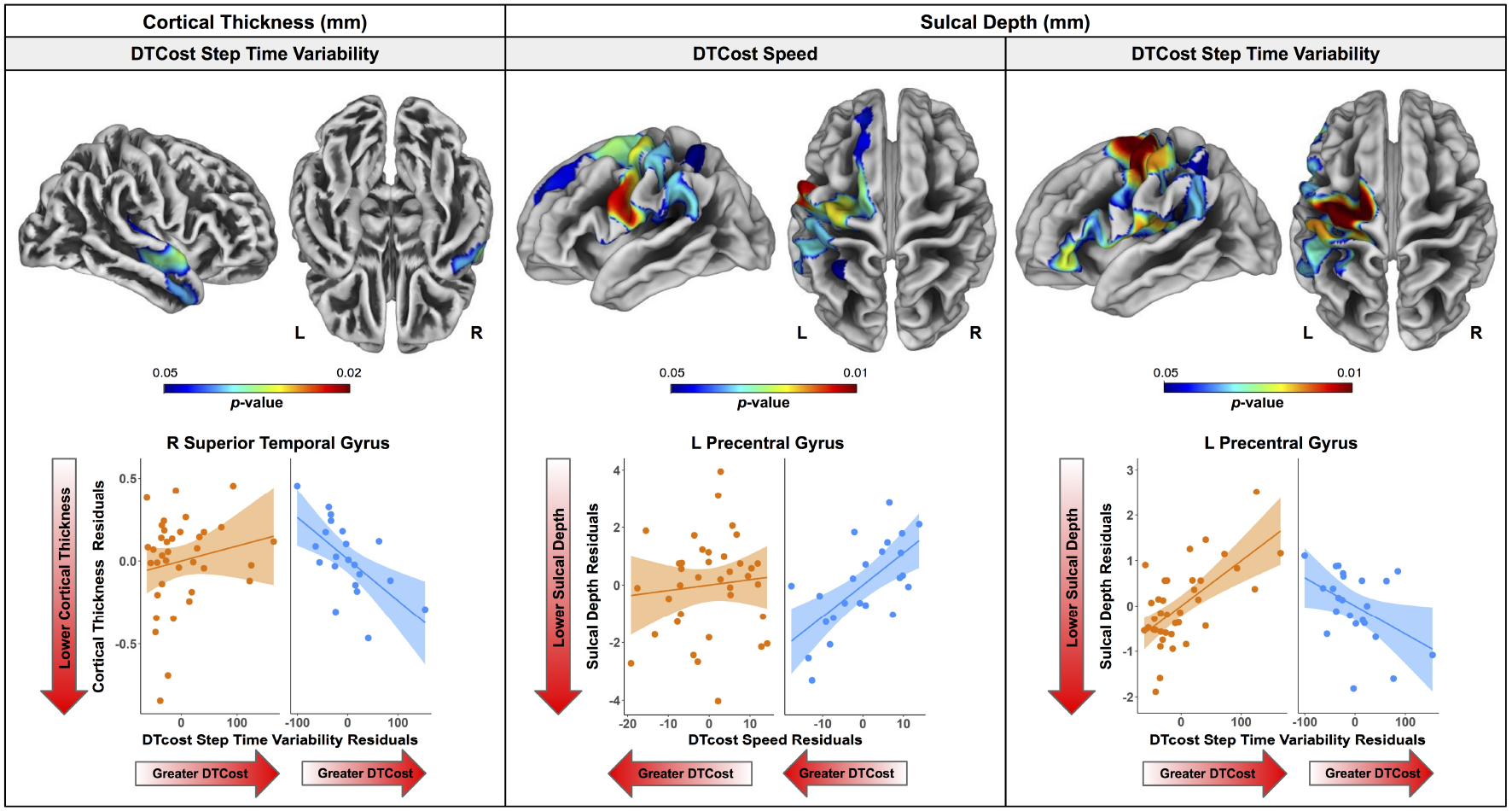
Age differences in the relationship of surface metrics with the DTcost of gait. Top. Regions showing statistically significant (*p*_*FWE− corr*_ < 0.05) age group differences in the relationship of cortical thickness (left) and sulcal depth (middle, right) with the DTcost of gait speed and step time variability. Warmer colors indicate regions of greater age differences in brain-behavior correlations. Results are overlaid onto CAT12 standard space templates. L = left; R = right. Bottom. Surface values for the peak result coordinate for each model are plotted against DTcost of gait to illustrate examples of the relationships identified by the voxelwise statistical tests. The fit line and confidence interval shading are included only to aid visualization of these relationships. We plotted the residuals instead of the raw values here to adjust for the effects of the sex covariate included in each model.

There were age differences in the relationship between DTcost of gait speed and both ADt and RDt in portions of the left superior corona radiata involving the superior longitudinal fasciculus and corticospinal tract (Fig. 7; Table 4). For the older adults only, higher ADt and lower RDt in these regions was associated with greater slowing of gait speed from single to dual task conditions. Young adults showed no relationship between ADt or RDt in these regions and DTcost of gait speed. There were no statistically significant age group differences in the correlation of FAt or FW with the DTcost of gait speed.

For older adults only, larger lateral ventricular volume was associated with greater decreases in gait speed from single to dual task walking (Fig. 8; Table 5). There was no relationship between lateral ventricular volume and DTcost of gait speed for young adults. Older adult relationships between DTcost of gait speed with several other ROIs (i.e., thalamus gray matter volume (*p* = 0.025; *p*_*FDR−corr*_ = 0.172) and parahippocampal cortex volume (*p* = 0.045; *p*_*FDR−corr*_ = 0.208)) did not survive FDR correction. There were no other statistically significant interactions between age group and DTcost of gait speed for the remaining ROIs.

**Table 4.** Regions of age difference in the relationship of FW-corrected white matter microstructure with the DTcost of gait speed

| Region | TFCE Level |  | MNI Coordinates (mm) |  |  |
| --- | --- | --- | --- | --- | --- |
| | Extent ($k_E$) | $p_{FWE-corr}$ | X | Y | Z |
| ADt |  |  |  |  |  |
| L corona radiata (superior) / superior long. fasciculus | 204 | 0.026* | -24 | -7 | 34 |
| L corona radiata (superior) / corticospinal tract | – | 0.027* | -26 | -15 | 31 |
| L corona radiata (superior) / superior long. fasciculus | – | 0.045* | -26 | 1 | 27 |
| RDt |  |  |  |  |  |
| L corona radiata (superior) / superior long. fasciculus | 126 | 0.034* | -24 | -7 | 34 |
| L corona radiata (superior) / corticospinal tract | – | 0.035* | -26 | -15 | 30 |

*Note:* Here we list up to three local maxima separated by more than 8 mm per cluster for all clusters with size $k > 10$ voxels. The clusters were labeled using two atlases: the Johns Hopkins University (JHU) ICBM-DTI-82 White Matter Labels (listed first, to the left side of the slash), and the JHU White Matter Tractography atlas within FSL (listed second, to the right side of the slash) (Hua et al., 2008; Wakana et al., 2007). The clusters were sorted by $p_{FWE-corr}$ value (from smallest to largest), then by cluster size (from largest to smallest). L = left; Long = longitudinal. \* $p_{FWE-corr} < 0.05$ .

**Table 5.**
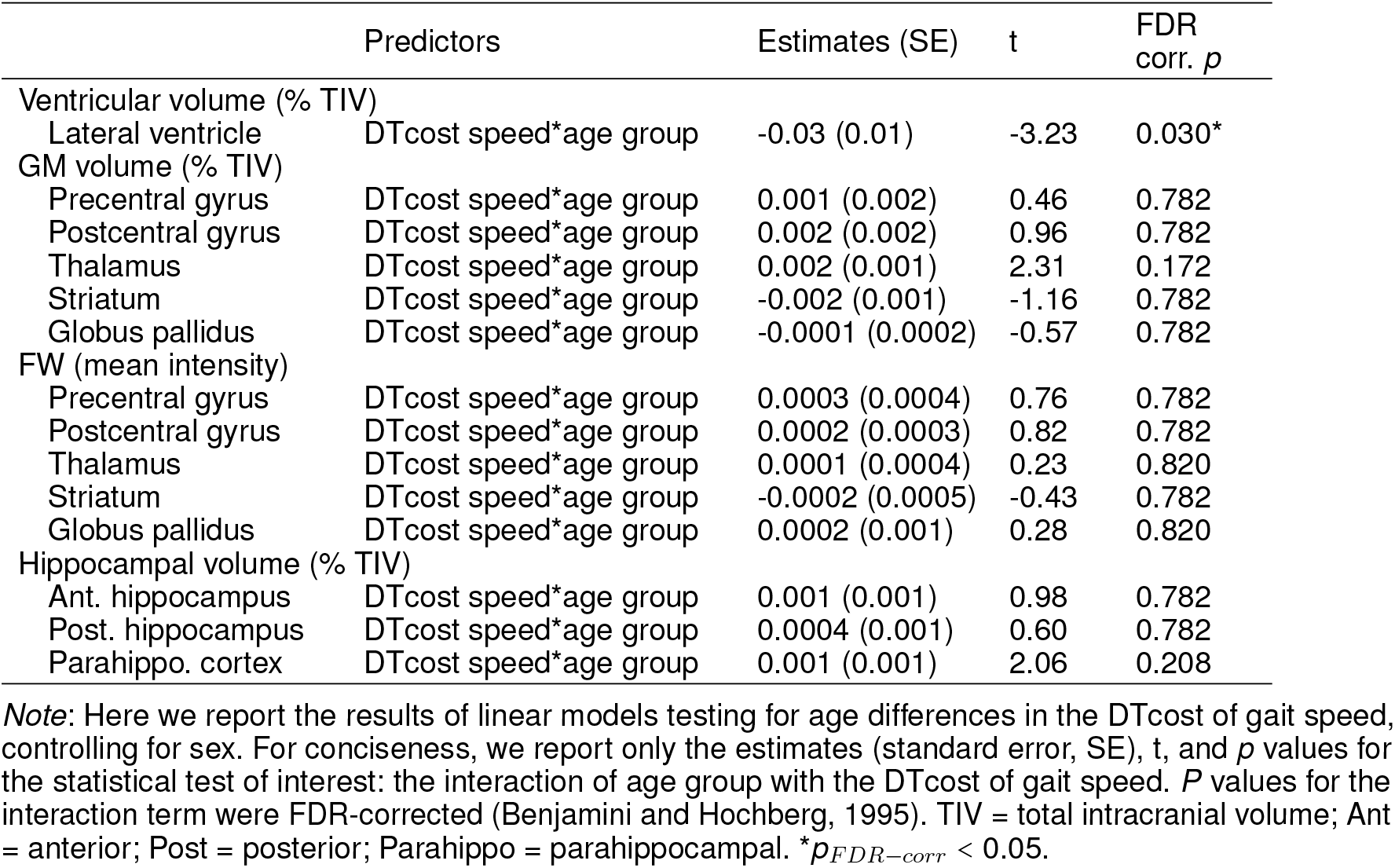
Regions of age difference in the relationship of structural ROIs with the DTcost of gait speed

|  | Predictors | Estimates (SE) | t | FDR<br>corr. *p* |
| --- | --- | --- | --- | --- |
| Ventricular volume (% TIV) |  |  |  |  |
| Lateral ventricle | DTcost speed*age group | -0.03 (0.01) | -3.23 | 0.030* |
| GM volume (% TIV) |  |  |  |  |
| Precentral gyrus | DTcost speed*age group | 0.001 (0.002) | 0.46 | 0.782 |
| Postcentral gyrus | DTcost speed*age group | 0.002 (0.002) | 0.96 | 0.782 |
| Thalamus | DTcost speed*age group | 0.002 (0.001) | 2.31 | 0.172 |
| Striatum | DTcost speed*age group | -0.002 (0.001) | -1.16 | 0.782 |
| Globus pallidus | DTcost speed*age group | -0.0001 (0.0002) | -0.57 | 0.782 |
| FW (mean intensity) |  |  |  |  |
| Precentral gyrus | DTcost speed*age group | 0.0003 (0.0004) | 0.76 | 0.782 |
| Postcentral gyrus | DTcost speed*age group | 0.0002 (0.0003) | 0.82 | 0.782 |
| Thalamus | DTcost speed*age group | 0.0001 (0.0004) | 0.23 | 0.820 |
| Striatum | DTcost speed*age group | -0.0002 (0.0005) | -0.43 | 0.782 |
| Globus pallidus | DTcost speed*age group | 0.0002 (0.001) | 0.28 | 0.820 |
| Hippocampal volume (% TIV) |  |  |  |  |
| Ant. hippocampus | DTcost speed*age group | 0.001 (0.001) | 0.98 | 0.782 |
| Post. hippocampus | DTcost speed*age group | 0.0004 (0.001) | 0.60 | 0.782 |
| Parahippo. cortex | DTcost speed*age group | 0.001 (0.001) | 2.06 | 0.208 |

*Note:* Here we report the results of linear models testing for age differences in the DTcost of gait speed, controlling for sex. For conciseness, we report only the estimates (standard error, SE), *t*, and *p* values for the statistical test of interest: the interaction of age group with the DTcost of gait speed. *P* values for the interaction term were FDR-corrected (Benjamini and Hochberg, 1995). TIV = total intracranial volume; Ant = anterior; Post = posterior; Parahippo = parahippocampal. \* $p_{FDR-corr} < 0.05$ .

**Figure 7.**
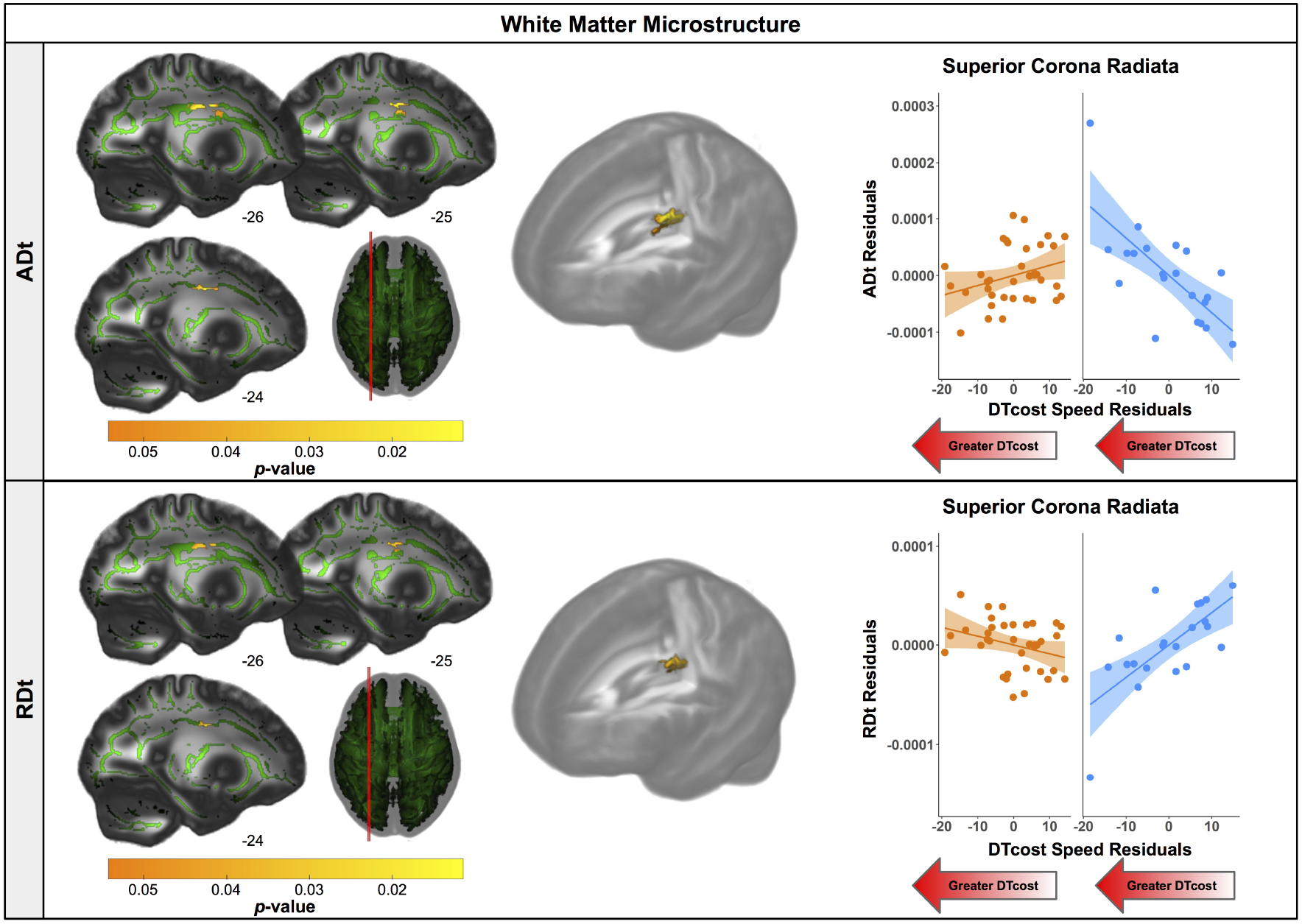
Age differences in the relationship of FW-corrected white matter microstructure with the DTcost of gait speed. Left. Regions showing statistically significant (*p*_*FWE− corr*_ < 0.05) age group differences in the relationship of ADt (top) and RDt (bottom) with the DTcost of gait speed. Warmer colors indicate regions of greater age differences. Results are shown on the FMRIB58 FA template with the group mean white matter skeleton (green) overlaid. Right. ADt and RDt values for the peak result coordinate for each model are plotted against the DTcost of gait speed to illustrate examples of the relationships identified by the voxelwise statistical tests. The fit line and confidence interval shading are included only to aid visualization of these relationships. We plotted the residuals instead of the raw values here to adjust for the effects of the sex covariate included in each model.

**Figure 8.**
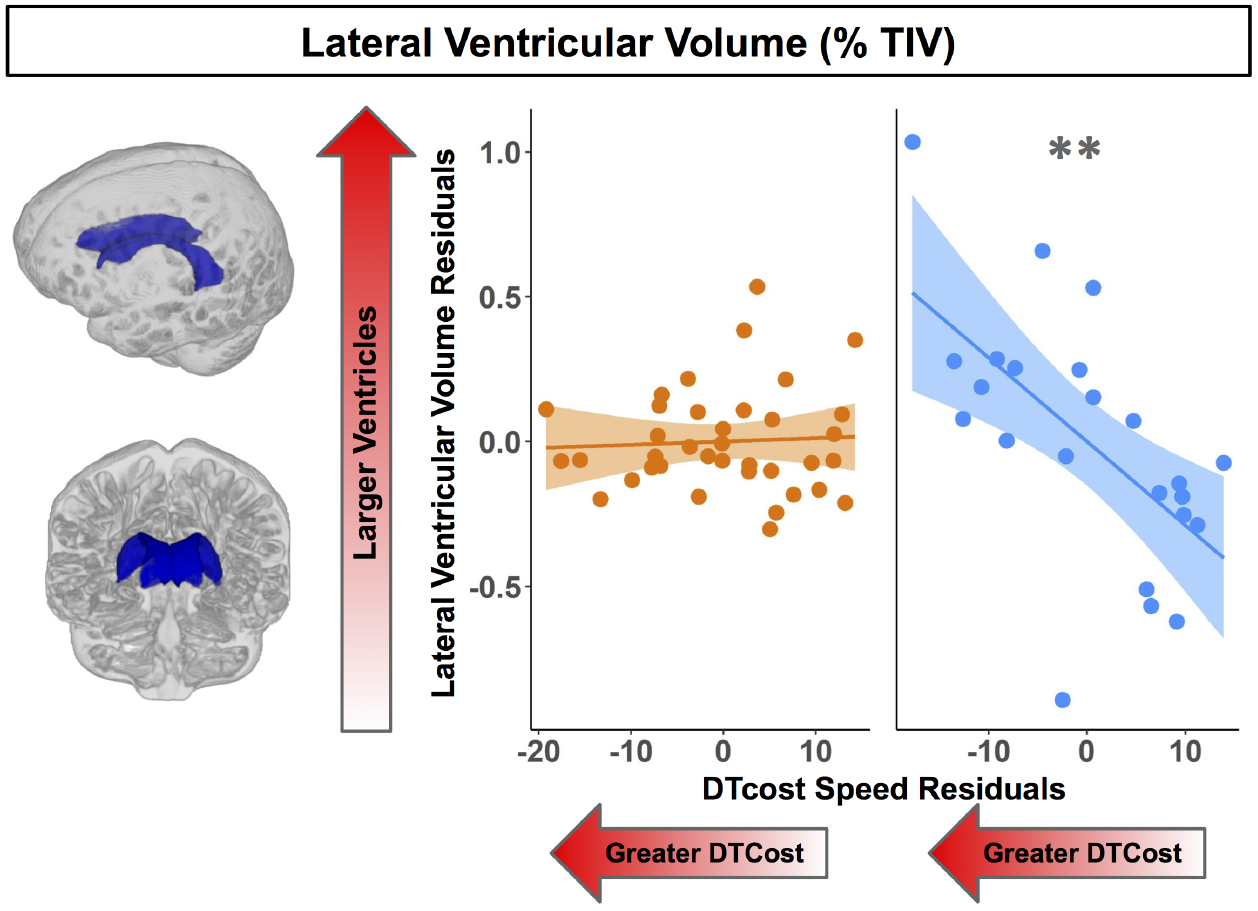
Age differences in the relationship of lateral ventricular volume with the DTcost of gait speed. Left. Here we depict the lateral ventricular volume mask for a single exemplar participant overlaid onto that participant’s native space cerebrospinal fluid segment. Right. Lateral ventricular volume residuals (expressed as a percentage of total intracranial volume) are plotted against the DTcost of gait speed. We plotted the residuals instead of the raw values here to adjust for the effects of the sex covariate included in the model. \*\**p*_*FDR−corr*_ < 0.01.

### 2.22 Age Differences in the Relationship of Brain Structure with the DTcost of Step Time Variability

There were no statistically significant age group by DTcost of step time variability interactions for gray matter or cerebellar volume. For older adults, thinner temporal lobe cortex was associated with greater DTcost of step time variability (Fig. 6; 6). That is, those older adults with the thinnest temporal cortex also showed the greatest increase in step time variability from single to dual task. Young adults showed a weak opposite relationship between temporal cortex thickness and the DTcost of step time variability. In addition, those older adults with shallower sulcal depth across the sensorimotor, supramarginal, insular, and superior frontal and parietal cortices also showed a greater DTcost of step time variability (Fig. 6; Table 3). Young adults showed a weak opposite relationship between sulcal depth in these regions and the DTcost of step time variability. There were no statistically significant age differences in the relationship of cortical complexity or gyrification index with the DTcost of step time variability.

**Table 6.** Regions of age difference in the correlation of cortical thickness with the DTcost of step time variability

| Region | Overlap of Atlas Region | TFCE Level |  |
| --- | --- | --- | --- |
| | | Extent ($k_E$) | $p_{FWE-corr}$ |
| DTcost of step time variability |  |  |  |
| R superior temporal gyrus | 68% | 790 | 0.032* |
| R middle temporal gyrus | 22% | — | — |
| R transverse temporal gyrus | 8% | — | — |

*Note:* Here we list all atlas regions from the Desikan-Killiany DK40 atlas (Desikan et al., 2006) that overlapped by 5% or more with the resulting cluster. We do not list volumetric (e.g., MNI space) coordinates in this table because volumetric coordinates cannot be mapped directly onto cortical surfaces. R = right.
\* $p_{FWE-corr} < 0.05$ .

There were no statistically significant age differences in the relationship between the DTcost of step time variability and FW-corrected white matter microstructure. Greater DTcost of step time variability was associated with lower parahippocampal cortex volume for the older adults, though this relationship did not survive FDR correction (*p* = 0.039; *p*_*FDR−corr*_ = 0.433). There were no statistically significant interactions between age group and the DTcost of step time variability for the remaining ROIs (Table S2).

### 2.23 Multiple Regression to Identify the Best Predictors of DTcost of Gait in Older Adults

For the DTcost of gait speed full model, we entered each participant’s left precentral gyrus sulcal depth and superior longitudinal fasciculus ADt and RDt (extracted from the peak region resulting from each voxelwise model). We also entered lateral ventricular volume (expressed as a percentage of total intracranial volume) and sex. The stepwise regression returned a model containing only sulcal depth, ADt, and sex, indicating that the combination of these three variables best predicts the DTcost of gait speed for older adults (Table 7).

**Table 7.**
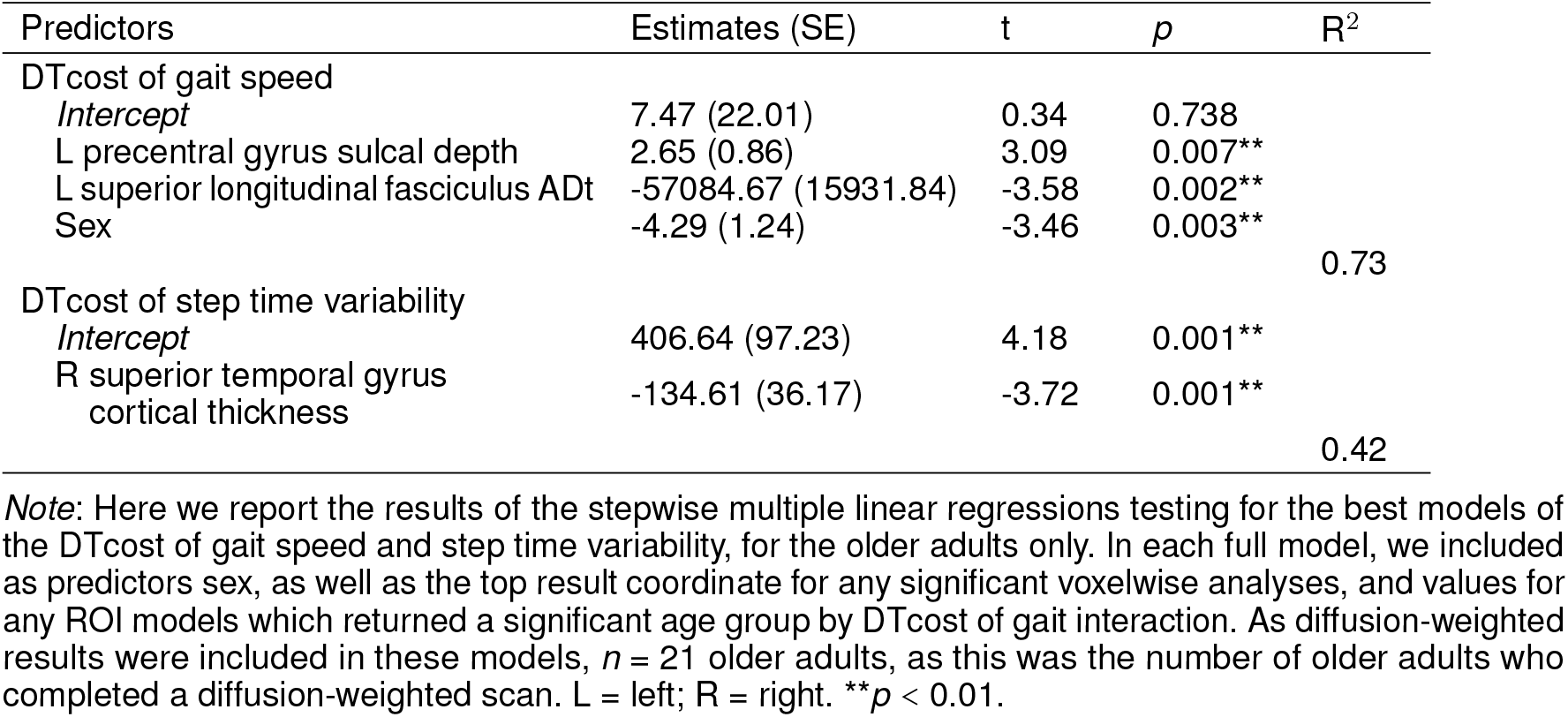
Stepwise multiple regression results for the best models of DTcost of gait in older adults

For the DTcost of step time variability full model, we entered each participant’s right superior temporal gyrus cortical thickness and left precentral gyrus sulcal depth, as well as sex. The stepwise regression returned a model containing only cortical thickness, indicating that this surface metric best predicts the DTcost of step time variability for older adults (Table 7).

## 3 DISCUSSION

We examined a comprehensive set of structural MRI metrics in relation to dual task walking in older adults. We identified widespread brain atrophy for older adults; across imaging modalities, we found the most prominent age-related atrophy in brain regions related to sensorimotor processing. Moreover, though the DTcost of gait speed and variability did not differ by age group, we identified multiple age differences in the relationship between brain structure and DTcost of gait. These age differences occurred both in regional metrics such as the temporal cortices and white matter tracts involved in motor control, and also for more general markers of brain atrophy, such as the lateral ventricles. We selected dual task walking performance as our outcome metric, as it is more predictive of falls in aging than single task walking (Ayers et al., 2014; Gillain et al., 2019; Halliday et al., 2018; Johansson et al., 2016; Verghese et al., 2017) and more related to real-world mobility (Hillel et al., 2019). Together, these results provide greater scientific understanding of the structural correlates of dual task walking in aging and highlight potential targets for future mobility interventions.

### 3.1 No Age Differences in the DTcost of Gait

Gait speed slowed, gait variability increased, and total number of subtraction problems attempted decreased between the single and dual task conditions. However, there were no age differences in the DTcost of gait speed, step time variability, or serial subtraction performance. That is, older adults did not exhibit a disproportionately larger decrease in gait speed or increase in gait variability between the NW and WWT conditions. Older adults also did not exhibit a disproportionately larger decrease in the total number of subtraction problems attempted between the seated and WWT conditions. While previous literature has mostly reported larger DTcosts to gait in older adults (e.g., for review see Al-Yahya et al., 2011; Beurskens and Bock, 2012), other previous work has found no age differences in the DTcost of gait speed (Holtzer et al., 2011). Moreover, much of this prior work has focused on comparisons of aging with pathologies such as cognitive impairment (Montero-Odasso et al., 2012; Pettersson et al., 2007), rather than comparisons of young and older adults. In our sample of relatively high-functioning older adults, the lack of group differences in the DTcost of gait and subtraction performance is perhaps unsurprising. Of note, we do believe that our cognitive task (serial 7s) was sufficiently difficult to divide attention between walking and talking for both age groups, as our task was more difficult than other common paradigms, such as reciting alternate letters of the alphabet (Ayers et al., 2014; Tripathi et al., 2019; Verghese et al., 2007). This lack of group differences in behavioral performance then frames our brain structure analyses to probe the neural correlates of preservation of function in aging. Thus, we can explore the neural correlates that might underlie compensation for normal brain aging and permit successful maintenance of dual task walking abilities into older age.

### 3.2 Age Differences in Brain Structure

#### 3.2.1 Gray matter volume, cerebellar volume, and cortical thickness

Overall, we found evidence of widespread brain atrophy for older compared with young adults. This observation is well in line with previous literature, which has similarly identified widespread age differences in brain gray matter volume (e.g., Lemaitre et al., 2012; Raz et al., 2010; Storsve et al., 2014), cerebellar volume (e.g., Bernard et al., 2015; Han et al., 2020; Koppelmans et al., 2017; Raz et al., 2010), and cortical thickness (e.g., Fjell et al., 2009b; Lemaitre et al., 2012; Salat et al., 2004; Storsve et al., 2014; Thambisetty et al., 2010; van Velsen et al., 2013). Many reports suggest that age-related atrophy occurs disproportionately in the frontal cortices (e.g., Fjell et al., 2009a; Lemaitre et al., 2012; Salat et al., 2004; Thambisetty et al., 2010). However, our finding of the most prominent age differences in gray matter volume and thickness of the sensorimotor cortices (and comparatively less age difference in the frontal cortices) fits with recent work which identified the greatest age differences (gray and white matter atrophy, demyelination, FW, and iron reduction) within the sensorimotor cortices in a large (*n* = 966) sample of middle- to older-aged adults (Taubert et al., 2020). Taubert and colleagues suggested that the particular age differences in sensorimotor cortex structure could be either a cause or an effect of age-related impairments to motor control (Papegaaij et al., 2014; Taubert et al., 2020).

#### 3.2.2 Additional surface metrics

While previous reports indicate that patterns of cortical thinning with aging largely mirror age-related changes in gray matter volume, the effects of aging on the other surface metrics studied here (i.e., sulcal depth, cortical complexity, and gyrification index) are not as well characterized. A couple of prior reports have indicated that, with aging, sulci become wider and shallower (Jin et al., 2018; Rettmann et al., 2006), and the cortex becomes less complex (Madan and Kensinger, 2016), with lower gyrification indices (Cao et al., 2017; Hogstrom et al., 2013; Lamballais et al., 2020; Madan, 2021; Madan and Kensinger, 2018). Our findings fit with these patterns, although across each of these metrics, we found the most prominent age differences within the lateral sulcus, whereas some previous work identified the largest age differences in other regions such as the central sulcus (cortical thickness; Rettmann et al., 2006), parietal lobe (sulcal depth; Jin et al., 2018), and frontal lobe (cortical complexity; Madan and Kensinger, 2016; and gyrification index; Lamballais et al., 2020). Methodological discrepancies might explain these differences; for instance, Jin et al. (2018) reported sulcal depth differences in middle versus older aged adults, rather than young compared with older adults.

#### 3.2.3 FW-corrected white matter microstructure

Only one previous study has directly compared FW corrected white matter microstructure between healthy young and older adults (Chad et al., 2018), despite that FW-corrected diffusion metrics have significantly higher test-retest reliability than conventional diffusion-weighted metrics (Albi et al., 2017), and that FW correction allows for separation of atrophy effects (i.e., increased extracellular fluid) from changes to the structure of the remaining white matter. Our findings here of age differences in FW-corrected white matter microstructure largely mirror those of Chad et al. (2018). As anticipated, we found lower FAt and ADt, paired with higher RDt and FW across almost the entire white matter skeleton. This pattern fits with previous literature examining FW-uncorrected white matter as well: prominent declines in FA, typically interpreted as decreased white matter microstructural organization and integrity (Bennett et al., 2010; Sexton et al., 2014) although also reflective of crossing fiber integrity (Chad et al., 2018), decreases in AD, interpreted as accumulation of debris or metabolic damage with age (Madden et al., 2012; Pierpaoli et al., 2001; Song et al., 2003), and increases in RD, interpreted as decreased myelin integrity or demyelination (Madden et al., 2012; Song et al., 2002, 2005).

After applying the FW correction to our data, we found several areas of opposite age differences, quite similar to the results described by Chad et al. (2018). Specifically, we observed a seemingly paradoxical finding in portions of the superior corona radiata, corpus callosum (e.g., splenium), internal capsule, and thalamic radiations, in which FAt and ADt were higher and RDt was lower for the older compared with the young adults. In addition to the report by Chad et al. (2018), several large datasets of normal aging (examining FW-uncorrected white matter) also corroborate this finding (de Groot et al., 2016; Miller et al., 2016; Sexton et al., 2014). Previous interpretations of this increased FA include selective degeneration of non-dominant tracts paired with a relative sparing of the primary bundle at fiber crossings (Chad et al., 2018). In particular, in this region, the corona radiata, internal capsule, and corpus callosum all cross the corticospinal tract (Tuch et al., 2003). The diffusion tensors in these regions indicate that the corticospinal tract is the principal fiber (Chad et al., 2018); *bedpostx* tractography analyses by Chad et al. (2018) suggest that the superior longitudinal fasciculus crosses the corona radiata in this region, and that the thalamic radiations also cross the corticospinal tract in this region of the internal capsule. Thus, as the superior longitudinal fasciculus and thalamic radiations are thought to degenerate substantially with age (Cox et al., 2016), while the corticospinal tract is thought to be relatively spared in aging (Jang and Seo, 2015), it is likely that the selective degeneration of non-dominant fibers in these locations is driving this seemingly paradoxical finding in the older adults.

#### 3.2.4 Structural ROIs

We selected the ROIs used in this study because of their purported roles in mobility function (i.e., the sensorimotor cortices, basal ganglia, and hippocampus; Beauchet et al., 2015, 2019; Callisaya et al., 2013). We also examined the lateral ventricles as a more general metric of subcortical atrophy. As anticipated, almost all of these ROIs showed significant age differences (i.e., reduced gray matter volume, increased FW, and increased ventricular volume). This fits with the existing literature reporting ventricular expansion in older age (Carmichael et al., 2009; Fjell et al., 2009a). However, it is interesting to note that FW fractional volumes showed less pronounced age differences compared to gray matter volumes. This could indicate that microstructural FW does not change as markedly with normal aging, in comparison to macrostructural gray matter tissue. Comparison of FW fractional volumes to prior aging work is difficult, as most previous papers report increased subcortical (e.g., substantia nigra) FW in pathological aging (e.g., Parkinson’s disease) compared with controls (Guttuso et al., 2018; Yang et al., 2019), as opposed to reporting comparisons of healthy young and older adults.

### 3.3 Interaction of Age Group with the DTcost of Gait

#### 3.3.1 Gray matter and cerebellar volumes

We did not identify any statistically significant age group differences in the relationship between the DTcost of gait speed or variability and regional gray matter volume. While extensive previous literature has examined relationships of single task overground walking with gray matter and cerebellar volume (e.g., Beauchet et al., 2015; Callisaya et al., 2013; Demnitz et al., 2017; Dumurgier et al., 2012; Rosano et al., 2007), comparatively less work has examined such relationships with dual task walking (Allali et al., 2019; Lucas et al., 2019; Ross et al., 2021; Tripathi et al., 2019; Wagshul et al., 2019). Further, these studies had methodological differences from our work (e.g., they used an alphabet task instead of serial 7s as the cognitive task). Moreover, it could be that we did not identify gray matter volume associations with the DTcost of gait because other measures (e.g., surface-based morphometry metrics) may provide a more sensitive correlate of behavior as compared with volume metrics. Surface-based metrics have been found to have several advantages over volume-based metrics (Hutton et al., 2009; Lemaitre et al., 2012; Winkler et al., 2010), including more accurate spatial registration (Desai et al., 2005), sensitivity to surface folding, and independence from head size (Gaser and Kurth, 2017).

#### 3.3.2 Surface metrics

We identified several age differences in brain-behavior relationships for two surface metrics: cortical thickness and sulcal depth. Only a few previous studies have examined relationships between cortical thickness and dual task walking in aging (Maidan et al., 2021; Ross et al., 2021), and, to our knowledge, no prior literature has examined sulcal depth in relation to dual task walking in aging. In the present work, we identified a relationship between thinner temporal cortex and greater increases in step time variability from single to dual task walking for older adults. Interestingly, the superior, middle, and transverse temporal gyri where we identified this result have functions in visual perception (Ishai et al., 1999; Miyashita, 1993), multimodal sensory integration (Downar et al., 2000; Mesulam, 1998), and spatial navigation (Howard et al., 2005). Given these functional roles, it is plausible that these regions of the temporal cortex would play a role in gait control.

Moreover, this region of temporal cortex is not one in which we found prominent age-related cortical thinning. Thus, it is possible that this temporal region plays a compensatory role in aging, to compensate for the substantial cortical thinning with aging that we identified in classical sensorimotor brain regions, such as the pre- and postcentral gyri. This notion fits with the hypothesis of neural inefficiency in aging (Fettrow et al., 2021b; Zahodne and Reuter-Lorenz, 2019), which suggests that, when neural resources become limited (as with age-related atrophy of the sensorimotor cortices), different neural resources (e.g., in this case, the temporal cortices) are used to compensate and maintain performance (e.g., as seen in the lack of age differences in the DTcost of gait). This also results in a stronger relationship between temporal lobe structure and dual task walking, which only emerges in older age when these neural resources start to become limited. This interpretation fits with a recent report of an association between lower cortical thickness and greater increases in prefrontal oxygenation from single to dual task walking, with no effect on performance (Ross et al., 2021). The study authors suggested that older adults with the poorest neural resources (i.e., the thinnest cortex) also required the most compensation from alternative brain regions (i.e., the greatest increases in prefrontal oxygenation) to maintain performance. One caveat to this interpretation, however, is that hypotheses of neural compensation with aging were largely developed in relation to functional, not structural, MRI data—though our data appear to follow a similar pattern.

We also identified two relationships between sulcal depth in aging and greater DTcost of gait speed and variability for older adults. Similar to cortical thickness, these brain-behavior relationships did not fall within the prominent regions of age difference in sulcal depth (i.e., the bilateral temporal lobes and insula), and instead spanned the sensorimotor, supramarginal, superior frontal and parietal cortices. Thus, these sulcal depth findings could similarly represent an age-related compensation. That is, in compensation for shallowing of other cortical regions in aging, those who retained deeper sulci into older age were also able to maintain the best functional walking performance.

Of note, while young adults did not show a clear relationship between cortical thickness or sulcal depth and DTcost of gait speed, young adults did exhibit a relationship between greater sulcal depth and lower DTcost of step time variability (which is in the opposite direction of what we might expect). Greater step time variability is clearly related to negative outcomes for older adults, such as higher fall risk (Callisaya et al., 2011). However, the case is less clear for young adults (Beauchet et al., 2009; Moe-Nilssen et al., 2010). For instance, higher gait variability for younger adults can indicate more stable gait (Beauchet et al., 2009). Additionally, it could be that young adults were using a different strategy to complete the task.

#### 3.3.3 FW-corrected white matter microstructure

Several prior studies have linked lower white matter diffusivity metrics to poorer overground walking (e.g., Bruijn et al., 2014; Tian et al., 2016; Verlinden et al., 2016) and dual task walking in older adults (e.g., Ghanavati et al., 2018). However, though one prior study identified relationships between FW-corrected white matter microstructure and cognition in normal aging (Gullett et al., 2020), to our knowledge, no previous work has examined how FW-corrected white matter microstructure relates to mobility in older adults.

We identified two relationships in which higher ADt and lower RDt were associated with worse dual task performance, i.e., greater slowing of gait speed from single to dual task conditions. This is perhaps the opposite pattern from what one might expect, as lower ADt is often associated with accumulation of debris or metabolic damage (Madden et al., 2012; Pierpaoli et al., 2001; Song et al., 2003), and higher RDt is interpreted as decreased myelin integrity or demyelination (Madden et al., 2012; Song et al., 2002, 2005). However, this result occurred in the superior corona radiata, where older adults had higher ADt and lower RDt than young adults (see Section 3.2.3). It could be that, in these white matter regions, the poorest performing older adults also have the greatest degeneration of crossing fibers, such as the superior longitudinal fasciculus crossing the corticospinal tract. As the superior longitudinal fasciculus is implicated in functions such as motor control, proprioception, and visuospatial attention and awareness (Amemiya and Naito, 2016; Rodríguez-Herreros et al., 2015; Shinoura et al., 2009; Spena et al., 2006), it is logical that deterioration of this pathway could negatively impact dual task walking in aging.

#### 3.3.4 Structural ROIs

We identified a relationship between larger lateral ventricular volume and greater DTcost of gait speed for older but not younger adults. This fits with some previous work that has linked larger ventricular volume with higher gait variability (Annweiler et al., 2014) and slower gait speed (Camicioli et al., 1999) in older adults. However, it is surprising that we did not identify relationships between DTcost of gait and the remaining structural ROIs, as previous work has linked sensorimotor (Rosano et al., 2007), basal ganglia (Dumurgier et al., 2012), and hippocampal (Beauchet et al., 2015) volumes to gait in aging. Our results thus suggest that generalized atrophy of subcortical structures, as opposed to atrophy of a single subcortical structure, is a better correlate of dual task locomotor function in aging.

### 3.4 Best Models of DTcost of Gait in Aging

Across the multimodal neuroimaging markers examined, left precentral gyrus sulcal depth, left superior longitudinal fasciculus ADt, and sex were the best predictors of DTcost of gait speed for older adults, and right superior temporal gyrus cortical thickness represented the best predictor of DTcost of step time variability. Given the purported benefits of surface metrics over volumetric measures (Desai et al., 2005; Hutton et al., 2009; Lemaitre et al., 2012; Winkler et al., 2010), the inclusion of sulcal depth and cortical thickness in these final models is perhaps unsurprising. Further, by minimizing partial volume effects resulting from white matter atrophy with aging, FW-corrected measures should provide greater sensitivity than traditional diffusion metrics for detecting true microstructural effects in aging cohorts. Thus, it is also perhaps unsurprising that ADt in a region (superior longitudinal fasciculus) particularly affected by aging (Cox et al., 2016) was also a good predictor of DTcost of gait in aging. Females showed larger DTcosts of gait speed, though previous literature has only infrequently reported sex differences in dual task walking in older adults (e.g., Hollman et al., 2011b; MacAulay et al., 2014; Yogev-Seligmann et al., 2010), and findings were conflicting.

Despite these results, we would also like to note that these surface and white matter metrics are complicated measures and that, although these produced the best models of DTcost of gait, it is worth mentioning that lateral ventricular volume also represented a good predictor of DTcost of gait speed in aging. Ventricular volume can be extracted easily by applying automated algorithms to common *T*_1_-weighted MRI sequences, and provides a useful general metric of subcortical atrophy, which our data suggest contributes functionally to gait speed slowing in aging.

### 3.5 Limitations

Our cross-sectional approach precluded us from tracking concurrent changes in brain structure and mobility over time. Additionally, our statistical models focused on the interaction of age group with the DTcost of gait, in order to identify regions where the relationship between brain structure and DTcost of gait differed for young versus older adults. We did not test for regions where brain structure related to DTcost of gait in the same manner for each age group. Such models may have uncovered more brain-behavior relationships in classical motor control regions, such as pre- and postcentral gyrus and the cerebellum. However, this was not a focus of the present work. Instead, our primary goal was to understand what brain regions contributed differently to maintenance of dual task walking in older age, to probe age-related shifts in the cortical control of gait and potential compensatory processes. In addition, we did not test for relationships between brain structure and subtraction performance. Subtraction accuracy did not differ between single and dual task conditions (i.e., most DTcost scores were close to 0) and thus it would not have made sense to assess brain-behavior relationships in this case. The total number of subtraction problems attempted was lower for both age groups during single compared to dual task, though this difference was less pronounced compared to the gait metrics. Future work could test whether there are different brain structure-behavior relationships for the DTcost of serial subtraction speed compared to the DTcost of gait metrics.

### 3.6 Conclusions

In this multimodal neuroimaging study, we found widespread age-related atrophy across cortical, subcortical, and cerebellar regions, but particularly in regions related to sensorimotor processing (e.g., the pre- and postcentral gyri). We then identified potential compensatory relationships between better maintenance of brain structure in regions not classically associated with motor control (e.g., the temporal cortices) and preserved dual task walking abilities in older adults. This suggests a role for the temporal cortices in maintaining behavioral function in aging, particularly when other brain regions responsible for locomotor control (e.g., the sensorimotor cortex, basal ganglia, and cerebellum) may be largely atrophied. Additionally, we identified one relationship between less specific subcortical atrophy (i.e., larger lateral ventricles) and greater slowing during dual task walking in aging. As the global population quickly ages, and emerging evidence continues to relate mobility problems with pathologies such as cognitive decline (Dodge et al., 2012; Knapstad et al., 2019), it is becoming increasingly critical to understand the structural neural correlates of locomotor function in aging. Identifying such brain markers could help identify those at the greatest risk of mobility declines, as well as identify targets for future interventions to preserve mobility and prevent disability among older adults.

## AUTHOR CONTRIBUTIONS

KH led the initial study design, collected and preprocessed all of the neuroimaging and gait data, conducted all statistical analyses, created the figures and tables, and wrote the first draft of the manuscript. JG assisted with data collection, data processing, and manuscript preparation. OP and HR consulted on DWI preprocessing and contributed to manuscript preparation. CH consulted on the design and analysis of the gait assessments. RS oversaw project design and led the interpretation and discussion of the results. All authors participated in revision of the manuscript.

## ACKNOWLEDGMENTS

The authors wish to thank Aakash Anandjiwala, Pilar Alvarez Jerez, and Alexis Jennings-Coulibaly for their assistance in subject recruitment and data collection, as well as Sutton Richmond for his help in applying the signal drift correction to the diffusion-weighted data. The authors also wish to thank all of the participants who volunteered their time, as well as the McKnight Brain Institute MRI technologists, without whom this project would not have been possible.

## DATA AVAILABILITY STATEMENT

The raw data supporting the conclusions of this article will be made available by the authors, without undue reservation.

