## Supplemental Information for "Differential relationships between brain structure and dual task walking in young and older adults"

### Supplementary Material

#### 1 SUPPLEMENTAL METHODS FOR PROCESSING OF DIFFUSION-WEIGHTED IMAGES

Here we provide more specific details regarding the preprocessing of the diffusion-weighted images.

**Signal drift:** We corrected the diffusion-weighted images images for signal drift (Vos et al., 2017) using the ExploreDTI graphical toolbox (University Medical Center Utrecht, Netherlands, Version 4.8.6; `www.exploredti.com`; Leemans et al., 2009) in MATLAB (R2019b).

**Topup:** We used the FMRIB Software Library (FSL)'s processing tool `topup` to estimate the susceptibility-induced off-resonance field (Andersson et al., 2003). We entered a pair of $b_0$ images collected with reversed phase-encode blips (i.e., the first volume of the diffusion-weighted sequence with Anterior to Posterior encoding and one $b_0$ volume from the Posterior to Anterior sequence collected immediately before the diffusion-weighted sequence). This procedure yielded a single corrected field map for use in eddy current correction.

**Eddy:** We used FSL's `eddy_cuda` to simultaneously correct the data for eddy current-induced distortions and both inter- and intra-volume head movement (Andersson and Sotiropoulos, 2016). We entered the topup-calculated field map with the `--topup` flag. We used the `--repol` flag to remove slices classified as outliers (i.e., where signal has been lost due to subject movement during the diffusion encoding) and replace these slices with non-parametric predictions by the Gaussian Process (Andersson et al., 2016). We set `--slm=1`, which specifies the mathematical form for how the diffusion gradients cause eddy currents; this setting is recommended when data are sampled on the half sphere, which was the case here. We used the `--estimate-movement-by-susceptibility` flag, which provides additional corrections to account for the effects of head movement on the diffusion signal (Andersson et al., 2018). We set `--mporder=17` to perform slice-to-volume movement correction (Andersson et al., 2017), which is a novel (but computationally expensive) method that corrects for within-volume head movement. We set the temporal order of movement at 17, as FSL documentation recommends using the number of excitations divided by 4 (i.e., # of slices / 4, so 69/4 $\approx$ 17). During `eddy_cuda`, rotations applied to each volume during motion correction were also applied to the corresponding *b* vectors. We then plotted each subject's volume-wise root mean square displacement provided by `eddy_cuda`. We considered a volume to be an outlier if its displacement was greater than 1 mm relative to the previous volume. Eight young and four older adults had one or more outlier volumes removed (young adults: 2-8 volumes removed per subject; older adults: 1-3 volumes removed per subject). Outlier volumes were removed from the eddy corrected image, as well as from the *b* value and rotated *b* vector matrices.

**Free-water (FW) Correction:** We implemented a custom FW algorithm (Pasternak et al., 2009) using MATLAB (R2019b). This algorithm estimates FW volume by fitting a bitensor model at each voxel of the preprocessed DWI image (Pasternak et al., 2009). The bitensor model consists of: 1) a tissue compartment, i.e., the diffusion indices (including FA, RD, and AD) of water molecules within white matter tissue; and 2) a FW compartment, reflecting the proportion of water molecules with unrestricted diffusion. FW fractional volumes range from 0 to 1; a fraction of 1 indicates that a voxel is filled with freely diffusing water molecules (e.g., as in the ventricles). The outputs of interest from this algorithm include a whole-brain FW map and FW-corrected whole-brain maps of white matter indices, denoted by subscript “t” to indicate that these metrics are based on the tissue compartment (FAt, RDt, and ADt).

**Tract-Based Spatial Statistics (TBSS):** We applied FSL’s tract-based spatial statistics (TBSS) processing steps to prepare the data for voxelwise analyses across participants (Smith et al., 2006). Benefits of TBSS include avoiding problems associated with suboptimal image registration between participants and eliminating the need for spatial smoothing. TBSS uses a carefully tuned nonlinear registration and projection onto an alignment-invariant tract representation (i.e., the mean FA skeleton); this process improves the sensitivity, objectivity, and interpretability of analyses of multi-subject diffusion studies.

First, we used `tbss_1_preproc` to erode the FA images slightly and zero the end slices (to remove likely outliers from the diffusion tensor fitting). Next, we used `tbss_2_reg` to calculate the warps to bring each subject’s FA data to a common space (i.e., the FMRIB58_FA 1 mm isotropic template) using the nonlinear registration tool FNIRT (Andersson et al., 2007b,a), which employs a b-spline representation of the registration warp field (Rueckert et al., 1999). We then used `tbss_3_postreg` to apply the warps calculated in step two, to calculate a mean FA image, and to thin this mean image to create a mean FA skeleton. This mean FA skeleton represented the centers of all tracts common to the whole group. Finally, we used `tbss_4_prestats` with a threshold of 0.2 to project each participant’s aligned FA data onto the group mean skeleton.

Lastly, we applied FSL’s `TBSS_non_FA` script to the additional whole-brain maps (i.e., the FW, FAt, RDt, and ADt maps). This applied the original nonlinear registration to these maps and projected the data onto the original mean FA skeleton (using the original FA data to find the projection vectors). Ultimately, these TBSS procedures resulted in skeletonized FW, FAt, ADt, and RDt maps in standard space for each participant. These were the maps that we entered in the group-level voxelwise statistical models described in the main text.

#### 2 SUPPLEMENTARY FIGURES AND TABLES

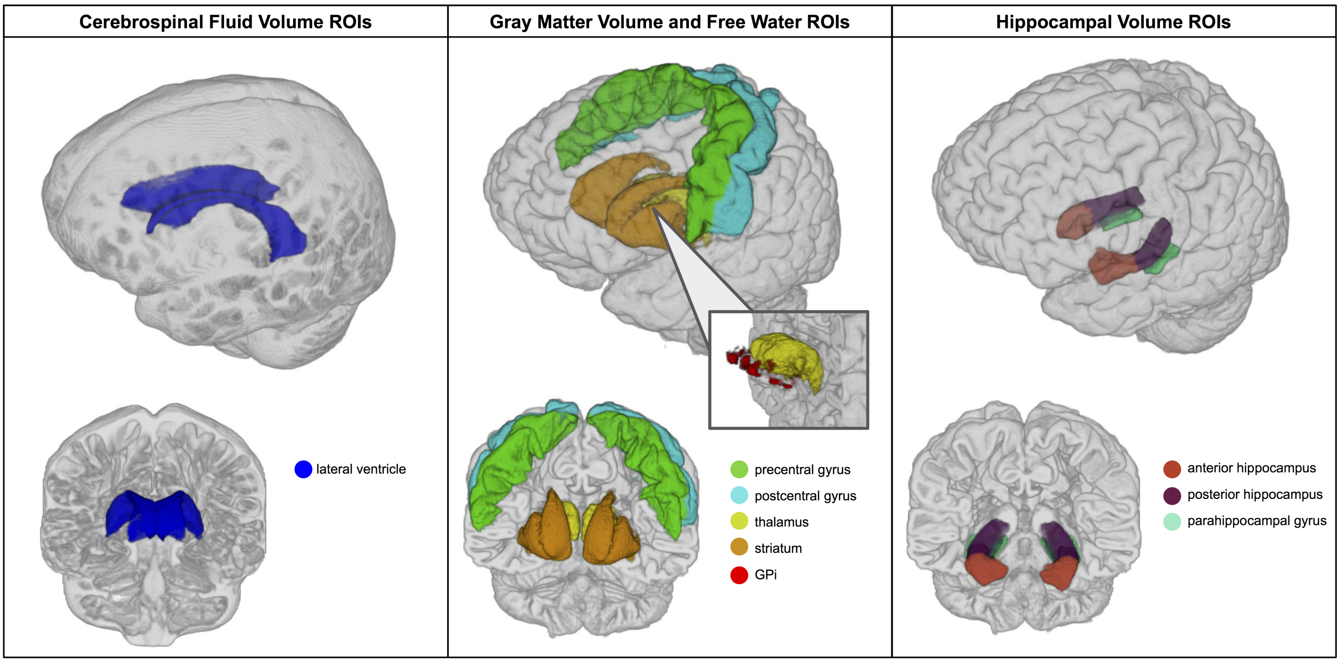


**Figure S1.** Structural ROIs. Here we depict ROI masks overlaid onto subject space cerebrospinal fluid (left) and gray matter (middle, right) segments for an exemplar young adult participant. In every case, we used the average of the left and right side ROI in our statistical analyses. Left. Lateral ventricle ROI masks. Middle. Five ROIs for which we extracted both gray matter volume and FW. ROIs are shown over the gray matter segment obtained from the $T_1$-weighted image. We do not depict here the subject space FW image from which the FW values were obtained. See Fig. S4 for illustrations of these ROIs overlaid onto the FW image. Right. Three hippocampal ROIs.

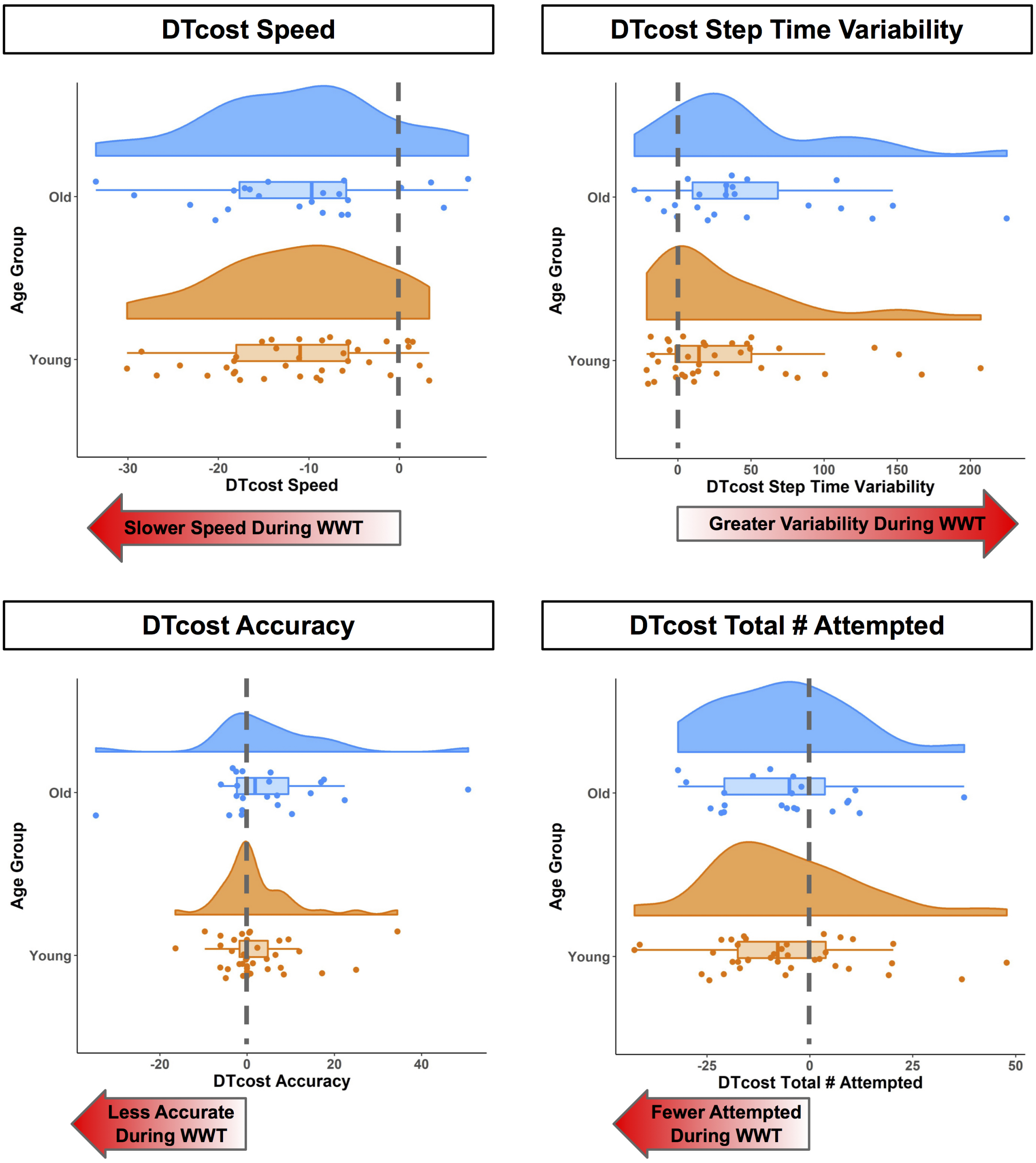


**Figure S2.** No age differences in the DTcost of gait and subtraction performance. The DTcost of gait and subtraction performance metrics is depicted for older (blue) and young (orange) adults. The red arrows indicate the direction of poorer performance during the dual compared to the single task conditions. There were no statistically significant age group differences in the DTcost of gait or serial subtraction performance. Gait speed was measured in m/s, step time variability was calculated as the standard deviation of step time, accuracy was calculated as the percent of subtractions problem answered correctly, and total number attempted was the total number of sutraction problems the participant attempted to answer.

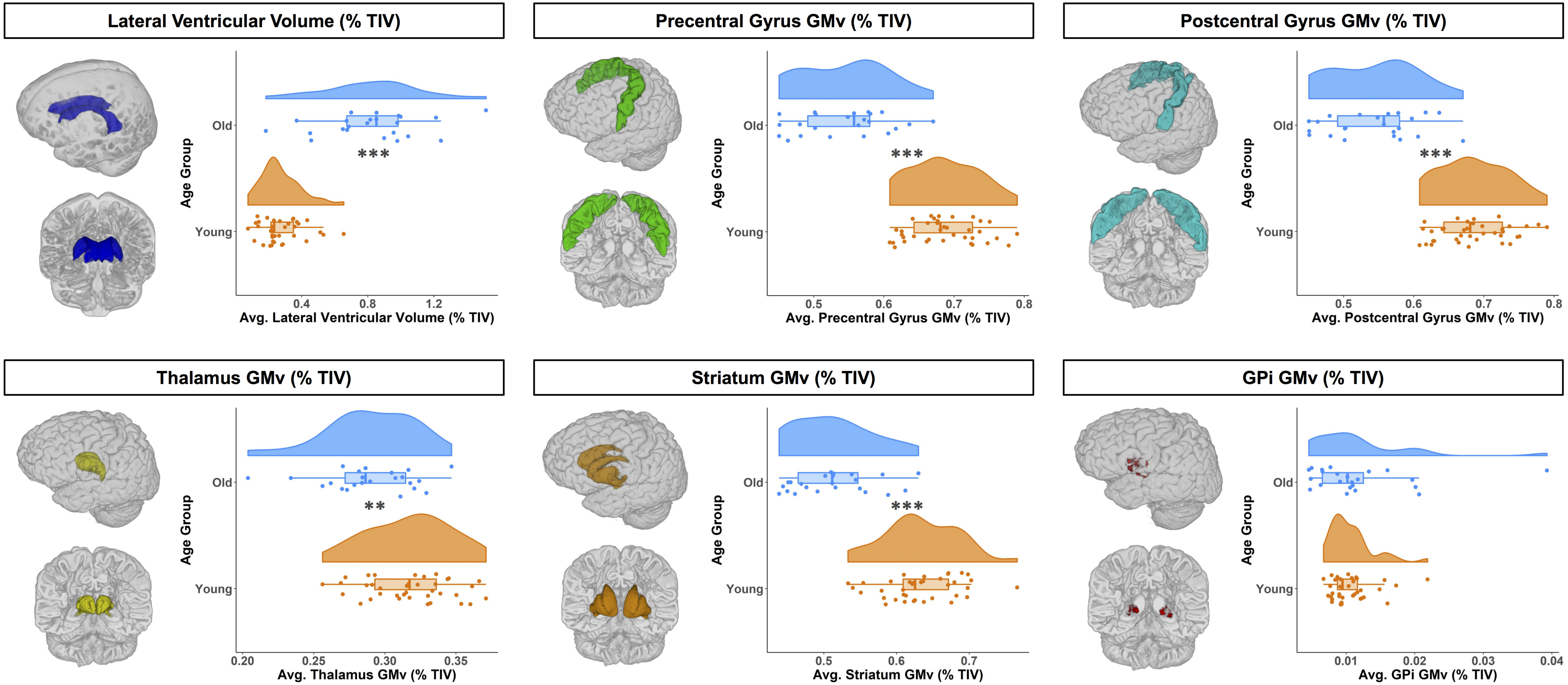


**Figure S3.** Age differences in ventricular volume and gray matter volume ROIs. For illustrative purposes, on the left we depict each structural ROI mask for a single exemplar subject overlaid onto that subject's native space cerebrospinal fluid (lateral ventricles) or gray matter segment (all other ROIs). On the right, we depict ROI gray matter volume (expressed as a percentage of total intracranial volume) for young (orange) and older (blue) adults. Compared with young adults, older adults had larger ventricles, as well as less gray matter volume across all ROIs except for the globus pallidus (GPi). $^{**}p_{FDR-corr} < 0.01$, $^{***}p_{FDR-corr} < 0.001$.

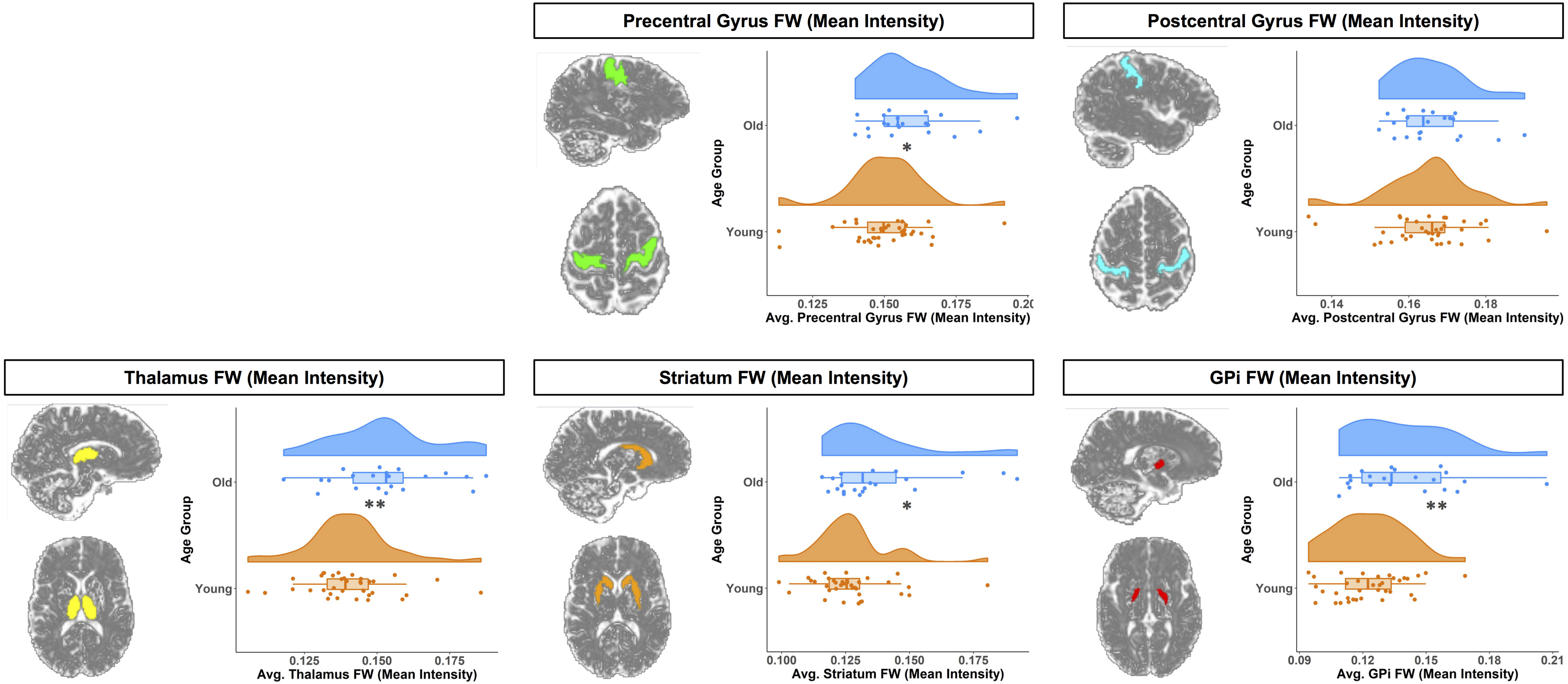


**Figure S4.** Age differences in FW ROIs. For illustrative purposes, on the left we depict each structural ROI mask for a single exemplar subject overlaid onto that subject's native space whole brain FW map. On the right, we depict ROI FW values (expressed as mean intensity) for young (orange) and older (blue) adults. Compared with young adults, older adults had higher FW values across all ROIs except for the postcentral gyrus. *$p_{FDR-corr} < 0.05$, **$p_{FDR-corr} < 0.01$.

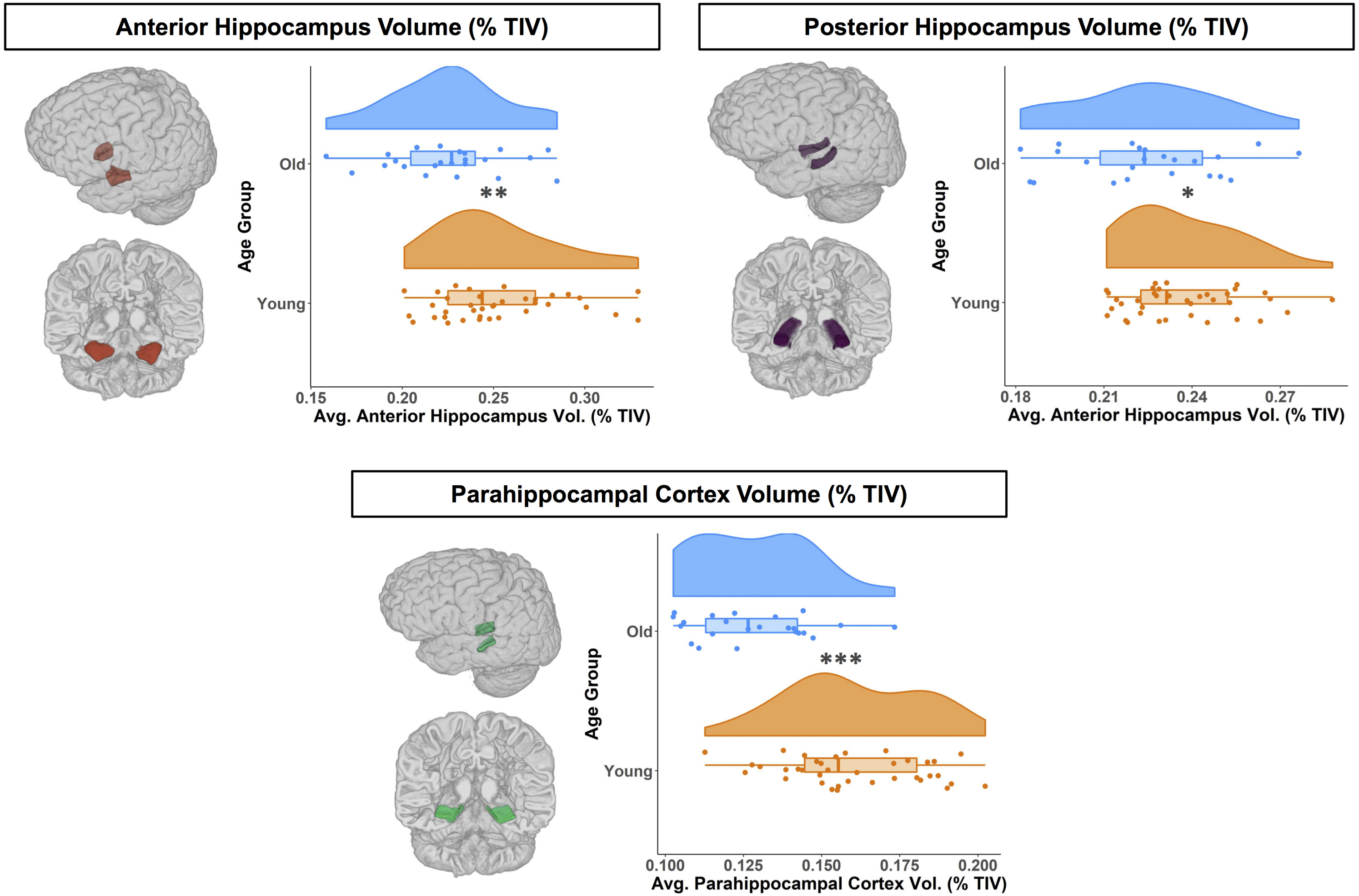


**Figure S5.** Age differences in hippocampal ROIs. For illustrative purposes, on the left we depict each hippocampal ROI mask for a single exemplar subject overlaid onto that subject's native space gray matter segment. On the right, we depict volume values (expressed as a percentage of total intracranial volume) for young (orange) and older (blue) adults. Compared with young adults, older adults had lower volume across all hippocampal ROIs. *$p_{FDR-corr} < 0.05$, **$p_{FDR-corr} < 0.01$, ***$p_{FDR-corr} < 0.001$.

**Table S1.** Age differences in structural ROIs

| Mean (SD) | | Predictors | Estimates (SE) | CI | t | FDR Corr. *p* | $R^2$ |
|---|---|---|---|---|---|---|---|
| Ventricular volume (% TIV) | | | | | | | |
| Lateral ventricle | | *(Intercept)* | 0.40 (0.05) | 0.30-0.50 | 7.97 | | |
| Young: 0.40 (0.18) | Old: 1.20 (0.44) | Age group *(Old)* | 0.80 (0.08) | 0.63-0.96 | 9.80 | $< 0.001$*** | |
| | | Sex *(Male)* | -0.02 (0.04) | -0.10-0.06 | -0.52 | 0.649 | |
| | | | | | | | 0.63 |
| GM volume (% TIV) | | | | | | | |
| Precentral gyrus | | *(Intercept)* | 0.86 (0.01) | 0.84-0.88 | 89.77 | | |
| Young: 0.86 (0.06) | Old: 0.64 (0.05) | Age group *(Old)* | -0.22 (0.02) | -0.25-(-0.19) | -14.07 | $< 0.001$*** | |
| | | Sex *(Male)* | 0.004 (0.01) | -0.01-0.02 | 0.66 | 0.649 | |
| | | | | | | | 0.78 |
| Postcentral gyrus | | *(Intercept)* | 0.69 (0.01) | 0.67-0.70 | 76.71 | | |
| Young: 0.69 (0.05) | Old: 0.54 (0.06) | Age group *(Old)* | -0.14 (0.01) | -0.17-(-0.11) | -9.93 | $< 0.001$*** | |
| | | Sex *(Male)* | 0.01 (0.01) | 0.00-0.03 | 1.66 | 0.159 | |
| | | | | | | | 0.64 |
| Thalamus | | *(Intercept)* | 0.32 (0.005) | 0.31-0.33 | 65.70 | | |
| Young: 0.32 (0.03) | Old: 0.29 (0.03) | Age group *(Old)* | -0.03 (0.01) | -0.04-(-0.01) | -3.59 | 0.002** | |
| | | Sex *(Male)* | 0.01 (0.004) | 0.00-0.02 | 2.60 | 0.042* | |
| | | | | | | | 0.26 |
| Striatum | | *(Intercept)* | 0.63 (0.01) | 0.62-0.65 | 75.10 | | |
| Young: 0.63 (0.05) | Old: 0.51 (0.06) | Age group *(Old)* | -0.12 (0.01) | -0.1-(-0.09) | -8.86 | $< 0.001$*** | |
| | | Sex *(Male)* | 0.01 (0.01) | 0.00-0.03 | 2.17 | 0.080 | |
| | | | | | | | 0.59 |
| Globus pallidus | | *(Intercept)* | 0.01 (0.001) | 0.01-0.01 | 12.03 | | |
| Young: 0.01 (0.003) | Old: 0.01 (0.01) | Age group *(Old)* | 0.001 (0.001) | 0.00-0.00 | 0.69 | 0.530 | |
| | | Sex *(Male)* | 0.0001 (0.001) | 0.00-0.00 | 0.17 | 0.866 | |
| | | | | | | | 0.01 |
| FW (mean intensity) | | | | | | | |
| Precentral gyrus | | *(Intercept)* | 0.15 (0.002) | 0.15-0.15 | 69.76 | | |
| Young: 0.15 (0.01) | Old: 0.16 (0.01) | Age group *(Old)* | 0.01 (0.004) | 0.00-0.02 | 2.41 | 0.028* | |
| | | Sex *(Male)* | 0.01 (0.002) | 0.00-0.01 | 3.46 | 0.013* | |
| | | | | | | | 0.25 |
| Postcentral gyrus | | *(Intercept)* | 0.16 (0.002) | 0.16-0.17 | 98.93 | | |
| Young: 0.16 (0.01) | Old: 0.17 (0.01) | Age group *(Old)* | 0.001 (0.003) | 0.00-0.01 | 0.43 | 0.667 | |
| | | Sex *(Male)* | 0.004 (0.001) | 0.00-0.01 | 3.20 | 0.013* | |
| | | | | | | | 0.16 |

Table S1. Continued

| Mean (SD) | | Predictors | Estimates (SE) | CI | t | FDR Corr. *p* | $R^2$ |
|---|---|---|---|---|---|---|---|
| FW (continued) | | | | | | | |
| Thalamus | | *(Intercept)* | 0.14 (0.003) | 0.13-0.15 | 55.65 | | |
| Young: 0.14 (0.01) | Old: 0.15 (0.02) | Age group *(Old)* | 0.01 (0.004) | 0.00-0.02 | 3.00 | 0.007** | |
| | | Sex *(Male)* | 0.01 (0.002) | 0.00-0.01 | 3.12 | 0.013* | |
| | | | | | | | 0.26 |
| Striatum | | *(Intercept)* | 0.13 (0.003) | 0.12-0.13 | 44.48 | | |
| Young: 0.13 (0.01) | Old: 0.14 (0.02) | Age group *(Old)* | 0.01 (0.005) | 0.00-0.02 | 2.33 | 0.030* | |
| | | Sex *(Male)* | 0.004 (0.002) | -0.00-0.01 | 1.98 | 0.101 | |
| | | | | | | | 0.15 |
| Globus pallidus | | *(Intercept)* | 0.12 (0.003) | 0.12-0.13 | 36.41 | | |
| Young: 0.12 (0.02) | Old: 0.14 (0.02) | Age group *(Old)* | 0.02 (0.01) | 0.00-0.03 | 2.90 | 0.008** | |
| | | Sex *(Male)* | 0.001 (0.003) | -0.00-0.01 | 0.55 | 0.649 | |
| | | | | | | | 0.14 |
| Hippocampal Volume (% TIV) | | | | | | | |
| Ant. hippocampus | | *(Intercept)* | 0.25 (0.01) | 0.24-0.26 | 47.64 | | |
| Young: 0.25 (0.03) | Old: 0.22 (0.03) | Age group *(Old)* | -0.03 (0.01) | -0.04-(0.01) | -3.18 | 0.005** | |
| | | Sex *(Male)* | -0.01 (0.004) | -0.02-0.00 | -1.94 | 0.101 | |
| | | | | | | | 0.20 |
| Post. hippocampus | | *(Intercept)* | 0.24 (0.004) | 0.23-0.24 | 65.85 | | |
| Young: 0.24 (0.02) | Old: 0.22 (0.03) | Age group *(Old)* | -0.01 (0.01) | -0.02-0.00 | -2.15 | 0.042* | |
| | | Sex *(Male)* | 0.003 (0.003) | -0.00-0.01 | 1.14 | 0.365 | |
| | | | | | | | 0.09 |
| Parahippo. cortex | | *(Intercept)* | 0.6 (0.003) | 0.15-0.17 | 48.17 | | |
| Young: 0.16 (0.02) | Old: 0.13 (0.02) | Age group *(Old)* | -0.03 (0.01) | -0.04-(-0.02) | -5.97 | $<$ 0.001*** | |
| | | Sex *(Male)* | 0.01 (0.003) | 0.00-0.01 | 2.25 | 0.080 | |
| | | | | | | | 0.42 |

*Note*: On the left, we report the mean (standard deviation) for the young and older age groups. On the right, we report the results of a linear model testing for age group differences in each ROI, controlling for sex. *P* values for the age group and sex predictors were FDR-corrected (Benjamini and Hochberg, 1995). SE = standard error; CI = 95% confidence interval; TIV = total intracranial volume; Ant = anterior; Post = posterior; Parahippo = parahippocampal. *$p_{FDR-corr} < 0.05$, **$p_{FDR-corr} < 0.01$, ***$p_{FDR-corr} < 0.001$.

**Table S2.** Regions of age difference in the relationship of structural ROIs with the DTcost of step time variability

| | Predictors | Estimates (SE) | t | FDR corr. *p* |
|---|---|---|---|---|
| Ventricular volume (% TIV) | | | | |
| Lateral ventricle | DTcost variability*age group | 0.003 (0.001) | 1.91 | 0.433 |
| GM volume (% TIV) | | | | |
| Precentral gyrus | DTcost variability*age group | -0.0004 (0.0003) | -1.40 | 0.553 |
| Postcentral gyrus | DTcost variability*age group | -0.0001 (0.0003) | -0.32 | 0.886 |
| Thalamus | DTcost variability*age group | -0.0002 (0.0001) | -1.61 | 0.524 |
| Striatum | DTcost variability*age group | 0.0001 (0.0002) | 0.43 | 0.886 |
| Globus pallidus | DTcost variability*age group | 0.00001 (0.00003) | 0.31 | 0.886 |
| FW (mean intensity) | | | | |
| Precentral gyrus | DTcost variability*age group | 0.00001 (0.0001) | 0.18 | 0.907 |
| Postcentral gyrus | DTcost variability*age group | -0.00002 (0.00005) | -0.51 | 0.886 |
| Thalamus | DTcost variability*age group | -0.00001 (0.0001) | -0.12 | 0.907 |
| Striatum | DTcost variability*age group | 0.0001 (0.0001) | 1.31 | 0.553 |
| Globus pallidus | DTcost variability*age group | -0.00004 (0.0001) | -0.41 | 0.886 |
| Hippocampal volume (% TIV) | | | | |
| Ant. hippocampus | DTcost variability*age group | -0.0001 (0.0001) | -0.97 | 0.780 |
| Post. hippocampus | DTcost variability*age group | -0.0001 (0.0001) | -0.86 | 0.786 |
| Parahippo. cortex | DTcost variability*age group | -0.0002 (0.0001) | -2.11 | 0.433 |

*Note*: Here we report the results of linear models testing for age differences in the DTcost of step time variability, controlling for sex. For conciseness, we report only the estimates (standard error, SE), t, and *p* values for the statistical test of interest: the interaction of age group with the DTcost of step time variability. *P* values for the interaction term were FDR-corrected (Benjamini and Hochberg, 1995). TIV = total intracranial volume; Ant = anterior; Post = posterior; Parahippo = parahippocampal. *$p_{FDR-corr} < 0.05$.
